# Protease suppression by native I9 inhibitor improves recombinant protein production in *Trichoderma reesei*

**DOI:** 10.64898/2026.05.05.722857

**Authors:** Mari Valkonen, Mykhailo Girych, Sami Havukainen, Emmi Sveholm, Jens C. Luoto, Georg Schmidt, Norman Adlung, Antti Aalto, O. H. Samuli Ollila, Hans Mattila, Tanja Paasela

**Author notes:** To whom correspondence should be addressed., Classification: Biological sciences; Applied Biological Sciences. These authors contributed equally to this work (Mari Valkonen, Mykhailo Girych). Authors have no financial or personal competing interest in connection with the manuscript and acknowledge all funding sources supporting the work.

## Abstract

Microbes are powerful cell factories for making molecules that are difficult or impossible to produce by other means. Filamentous fungi such as *Trichoderma reesei* are superior hosts for recombinant protein production, yet secreted proteases often degrade target proteins, reducing yields and limiting process robustness. Preventing proteolysis without relying on expensive commercial inhibitors or compromising strain fitness has been a longstanding challenge in fungal biotechnology, particularly for scaling up production to industrially relevant levels.

Here, we report the identification of a native inhibitory protein from *T. reesei*, TrI9, and show that directing its secretion into the culture medium markedly reduces extracellular protease activity and enables production of the highly protease-sensitive spider silk–like protein CBM-AQ12-CBM, which could be used in high-performance biomaterials. Computational and *in vitro* analyses provide mechanistic insight, showing that TrI9 functions as a multi-target inhibitor of subtilisin-like proteases (SLPs), including SLP2, a protease that cannot be eliminated by gene deletion due to its crucial role in normal growth and development.

This TrI9-based strategy for protease mitigation presents a novel approach for strain improvement, protecting a wide range of protease-labile products beyond silk proteins during large-scale fermentation. Protecting sensitive proteins without added inhibitors offers a cost-effective alternative for scaling up protein production across multiple applications, from protein-based materials manufacturing to pharmaceuticals, as well as enzyme and food applications.

**Significance:** The discovery and characterization of an independently encoded I9 inhibitor (TrI9) in the filamentous fungus *Trichoderma reesei* reveals a new layer of biological regulation of subtilisin-like protease activity in fungi. We show that overexpressing and secreting TrI9 into the culture supernatant can suppress extracellular proteolysis, including activity from the essential SLP2 protease thereby overcoming a major barrier to production of protease-sensitive recombinant proteins without compromising strain fitness. For industrial biomanufacturing, this translates into higher effective titers, more consistent product integrity and reduced reliance on costly commercial protease inhibitors, hence, improving process robustness and economics. As a broadly applicable strain-and-process strategy, TrI9-enabled protease control strengthens *T. reesei* as a scalable platform for precision fermentation of diverse, otherwise hard-to-produce proteins.

## Introduction

Microbial manufacturing enables production of complex proteins including pharmaceuticals, food and materials from renewable and low-cost carbon sources. For some proteins with exceptional properties, such as spider silk, large-scale production is only feasible through microbial processes. *Trichoderma reesei* is a widely used industrial production host for many major enzyme manufacturers and precision fermentation companies due to its exceptional secretory capacity, which can exceed 100 g/L of total secreted proteins under optimized conditions (1). However, proteolytic degradation remains a major bottleneck for heterologous protein production, often resulting in substantially lower yields compared to native proteins hindering their industrial application. While deletion of problematic protease genes can reduce target degradation, this approach frequently leads to adverse phenotypic effects. Alternative strategies to reduce protease activities include bioprocess optimization, the use of chemical protease inhibitors, and the identification and deletion of transcriptional regulators controlling protease gene expression (reviewed in (2)). One example of the latter approach is the deletion of the protease regulatory factor PEA1 in *T. reesei* (3, 4).

Detailed proteomic analysis has revealed that *T. reesei* secretes a variety of proteases, including aspartic, serine, glutamic, trypsin-like, and metalloproteases, each contributing to protein degradation under varying growth conditions. The secreted proteases that are essential for nutrient acquisition through organic nitrogen recycling (5) lower the yields of heterologous proteins, as demonstrated in studies linking protease activity to reduced recombinant protein titres (6). To avoid target degradation, deletion of nine secreted proteases in *T. reesei* increased the production of human interferon alpha-2b (IFN α-2b) to 2.4 g/L (7). Nevertheless, supplementing the cultivation with protease inhibitors led to even higher IFN α-2b yields, suggesting that some residual protease activity persisted in these deletion strains (7).

Limitations to deletion approach are that some proteases cannot be removed from production hosts without negative phenotypes arising from their deletions (8, 9). Subtilisin-like protease 2 (SLP2) has been identified as a particularly problematic contributor to protein degradation in *T. reesei*. Its activity is consistently detected in culture supernatants (5, 8, 9) and is linked to degradation of recombinant products, yet *slp2* cannot be deleted without severe defects in growth and sporulation (7, 10). This combination makes SLP2 one of the key obstacles for producing protease-sensitive target proteins at industrially relevant titres. SLP2 and other subtilisin-like proteases are produced as inactive precursors, zymogens, that are proteolytically activated by cleavage of an N-terminal I9 inhibitor propeptide domain that also putatively assists in the correct folding of the protease (11-13). The N-terminal I9 inhibitor propeptides of subtilases are not the only members of the I9 family. Independent genes encoding for I9 subtilisin propeptide-like inhibitors have been discovered. The first ones identified were two similar inhibitors of the vacuolar proteinase B in *S. cerevisiae*, with PBI2 being the predominant form (14, 15). Other related small inhibitors PoIA1 and PoIA2 were later identified in *Pleurotus ostreatus* (16). In addition to fungi, I9 propeptide-like inhibitors have been reported in plants (17).

Here we present a protection strategy against proteolytic degradation based on computationally identified independent I9 inhibitor protein encoded by *trire2_122168* gene in *T. reesei*. We demonstrate this approach by overexpressing and secreting the inhibitor protein into the culture medium. Using computational modelling and *in vitro* analysis, we show that the inhibitor exhibits potent inhibitory activity against subtilisin-like proteases in *T. reesei*, and provide mechanistic insights into its mode of action. Secretion of TrI9 under industrially relevant bioreactor conditions enables production of the protease-sensitive spider silk protein CBM-AQ12-CBM while markedly reducing total extracellular protease activity, paving the way for improved yields across a wide range of protein targets and industrial production of spider silk proteins.

## Results

### CBM-AQ12-CBM is degraded by SLP2

Spider silk-like protein CBM-AQ12-CBM is very unstable in *T. reesei* culture supernatants, but its stability can be improved by chemically inhibiting serine-type proteases (Fig. S1). To improve the production levels of CBM-AQ12-CBM, additional proteases were selected to be deleted from the current 11 protease deletion background. SLP2 (*trire2_123244*) was selected as one of the protease candidate for further investigation, as it has been consistently identified as one of the most abundant subtilisin-like proteases in *T. reesei* culture supernatants and is a known contributor to the degradation of heterologous proteins (7, 8).

CBM-AQ12-CBM is tagged with N-terminal His-tag and C-terminal Strep II tag which allows us to follow its production levels using western blotting. Deletion of *slp2* improved substantially the stability of CBM-AQ12-CBM as the fragmentation pattern was not as evident in deletion strains compared to strains with functional SLP2 (Fig. S2c). However, deletion of *slp2* caused severe growth and sporulation defects as also reported in previous studies(8) underscoring the need for alternative strategies to mitigate its activity (Fig. S2b).

To investigate how SLP2 target the AQ12 silk domain, we computationally mapped its putative cleavage sites within the AQ12 sequence using an AlphaFold2-based fragment docking approach evaluated with the actifpTM confidence metric (18, 19) (see Methods). This analysis suggested that AQ12 may be susceptible to cleavage at the substrate motif:

> ···PGQQ↓GPGQ···

where ↓ indicates the predicted scissile bond. Each of the twelve tandem repeats comprising AQ12 contains two occurrences of this motif, suggesting up to 24 potential cleavage sites distributed along the full-length AQ12 domain, which would be consistent with the extensive proteolytic susceptibility of this substrate.

### Discovery of the subtilisin like protease inhibitor TrI9

As SLP2 and/or other serine proteases seem to degrade CBM-AQ12-CBM but the deletion strategy was not optimal, we hypothesized that production of the target could be enhanced by inhibiting serine protease activity. *Saccharomyces cerevisiae* produces inhibitor protein Pbi2 that has been reported to inhibit Prb1 (YscB), a *T. reesei* SLP2 homolog (15), which prompted us to examine the *T. reesei* genome for a similar inhibitory protein that could be co-expressed with CBM-AQ12-CBM to enhance its stability. By searching for a homologue of the yeast PBI2, we identified a novel I9 inhibitor encoding gene, *trire_122168,* in the *T. reesei* QM6a genome, and designated the encoded protein as TrI9. Although the amino acid sequence identity to yeast Pbi2 was only 28 %, FoldSeek structure-based analysis (20) confirmed the identification of a 151-residue I9 inhibitor protein encoded by *trire2_122168*. Despite the modest sequence identity, structural superposition revealed remarkable conservation of the overall fold between the *T. reesei* TrI9 protein and yeast PBI2 (Fig. S3).

To investigate whether TrI9 could function as an SLP2 inhibitor, we performed computational structural analyses of its predicted protease binding mode. AlphaFold3 modelling (21) of full-length SLP2 confirmed the presence of an N-terminal I9 autoinhibitory domain, consistent with the established mechanism by which subtilisin-like proteases maintain their zymogen state through propeptide-mediated inhibition during vacuolar trafficking (22, 23). The AlphaFold3 model of full-length SLP2 represents this inhibited zymogen complex, where the I9 autoinhibitory domain (residues 27–139) remains bound following autocatalytic cleavage at M136 (P1 position), with the connecting loop (residues 140–152) threading through the prime-side subsites of the substrate-binding cleft as the polypeptide chain continues into the catalytic domain (residues 153–474) (Fig.1, top). AlphaFold3 co-folding of the novel TrI9 inhibitor with the SLP2 catalytic domain (lacking its endogenous N-terminal inhibitor) revealed a binding mode strikingly similar to that of the native autoinhibitory domain. In both cases, the C-terminal regions of the inhibitors occupy the substrate-binding cleft and catalytic pocket, while the structured core of each inhibitor domain associates with the lateral surface of the catalytic domain (Fig. 1). Notably, whereas the SLP2 autoinhibitory domain engages subsites P9 through P1 and continues through the prime-side positions via the connecting loop, TrI9 terminates at its C-terminus at the P1 position (residue Gln151), acting as a pure competitive inhibitor without prime-side contacts (Fig. 1, bottom).

**Fig. 1.**
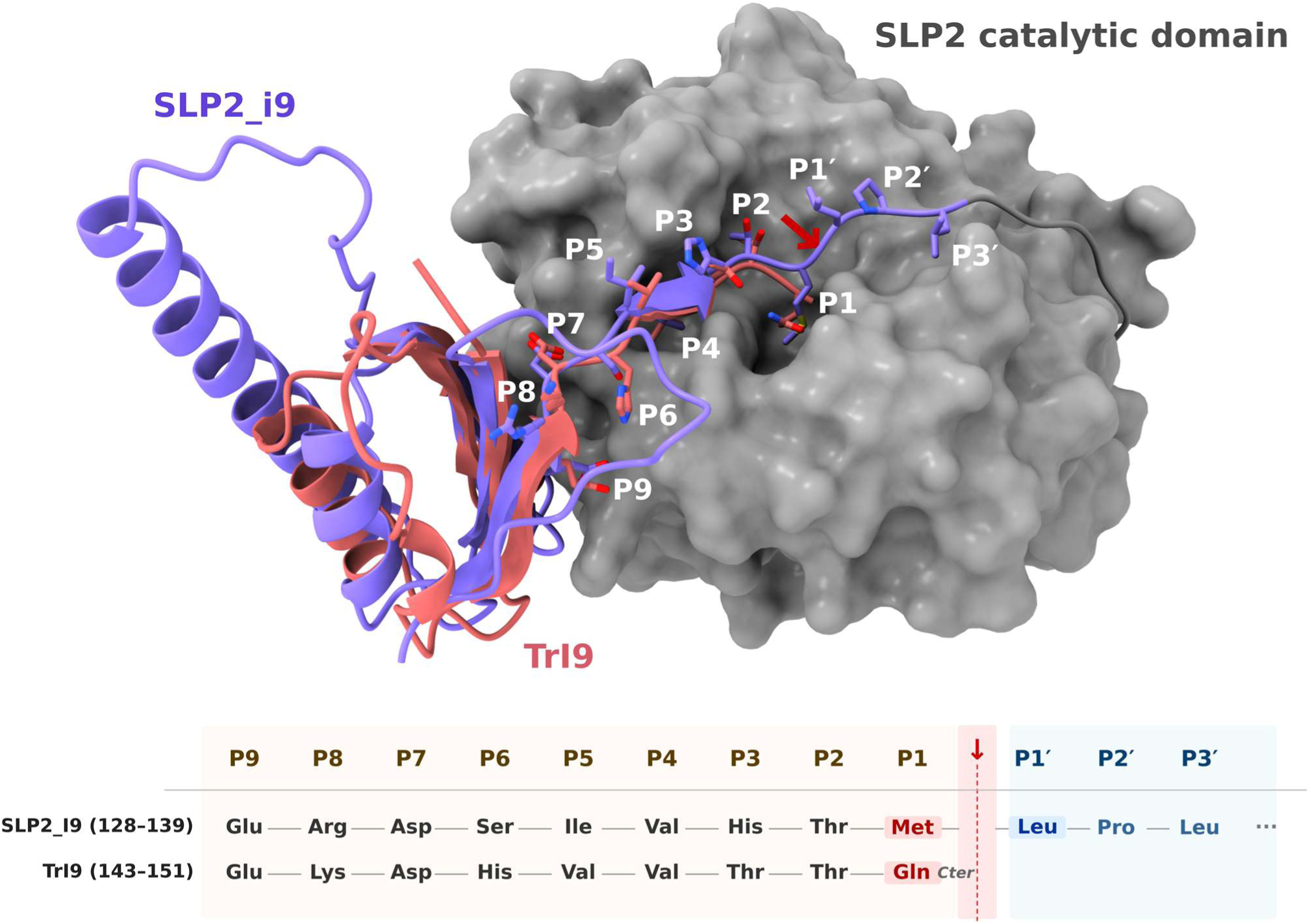
Structural basis of SLP2 inhibition by its autoinhibitory domain and the TrI9 inhibitor. (*Top*) AlphaFold3 models of the SLP2 **catalytic** domain (residues 153–474, gray surface) bound to SLP2_I9 (SLP2 autoinhibitory I9 domain, residues 27–139, purple cartoon) and TrI9 (residues 81–151, red cartoon), shown as an overlay. In both models, the inhibitor C-terminal loop occupies the substrate-binding cleft, while the structured core associates with the lateral surface of the catalytic domain. P-site residues (P9–P3′) are shown as sticks and labeled; the red arrow indicates the scissile bond. (*Bottom*) Substrate-like binding register of the C-terminal loop of each inhibitor: residue identities at positions P9–P3′ for SLP2_I9 (residues 128–139) and TrI9 (residues 143–151). The cleavage site (↓) separates the non-prime (P) from the prime (P′) positions.

Structural analysis of TrI9 reveals a bipartite architecture: residues 81–151 constitute the functional inhibitory domain that engages the SLP2 catalytic domain, whereas residues 1–80 are predicted to be intrinsically disordered and were therefore omitted from structural comparisons. Although sequence identity between the inhibitory domain of TrI9 (residues 81–151) and the SLP2 autoinhibitory domain (residues 27–136) is limited (23.9 %), their sequence similarity is considerably higher (53.5 % strict; 70.4 % inclusive), consistent with their shared structural and functional features. In particular, the C-terminal loops of both inhibitors, spanning residues 128–139 in SLP2_I9 and residues 143–151 in TrI9, show a conserved pattern of residue identities across the P9–P1 positions (Fig. 1, bottom), suggesting convergent adaptation to the same non-prime binding interface (24-26).

AlphaFold3 predictions for both the native SLP2 autoinhibitory domain and TrI9 binding to the SLP2 catalytic domain demonstrated high confidence metrics (SLP2_I9: interface PAE ∼7.1, min_ipSAE ∼0.78; TrI9: interface PAE ∼5.1, min_ipSAE ∼0.77) (25, 26), strongly supporting functional inhibitory interactions in both cases. Rosetta-based binding free energy calculations (24) yielded interface ΔG values of −91.84 and −80.89 REU for the native autoinhibitory domain and TrI9, respectively, indicating comparable predicted affinities with a moderate energetic advantage for the native autoinhibitory domain.

Intriguingly, extending this computational analysis to all nine annotated subtilisin-like proteases encoded in the *T. reesei* QM6a genome revealed that SLP2 may not be the sole target of TrI9. AlphaFold3 co-folding predictions and Rosetta affinity estimations, ranked by decreasing AF3 ipSAE_min interface confidence score, indicated favourable binding to the protease domains of SLP2 (*trire2_123244*), SLP3 (*trire2_123234*), SLP6 (*trire2_121495*), SLP5 (*trire2_64719*), and SLP8 *(trire2_58698*), with ipSAE_min values ranging from 0.709 to 0.769, mean interface PAE values below 5.1 Å, and Rosetta interface ΔG values between −71 and −86 REU. The remaining four SLPs, namely SLP10 (*trire2_35726*), SLP9 (trire2_60791), SLP4 (*trire2_109276*), and SLP1 *(trire2_51365*), yielded ipSAE_min values below 0.2 and mean interface PAE values exceeding 11 Å, indicating low confidence in the predicted binding geometry, yet in all cases TrI9 binding pose was predicted to be in the canonical inhibitory configuration (Fig. S4). Across all predicted complexes, the C-terminal loop of TrI9 was accommodated within the substrate-binding cleft of the respective protease, with the P1 residue positioned in the catalytic pocket. These findings suggest that TrI9 may function as a broad-spectrum inhibitor of multiple SLP family members.

### Synthetic TrI9 peptide inhibits the SLP2 protease activity *in vitro*

To verify the inhibitory activity of the TrI9 inhibitor against SLP2, we expressed and purified a recombinant SLP2. Production of full-length SLP2 was not successful. Expression of a truncated SLP2 variant (amino acids 15–469) in *T. reesei* with CBH1 signal sequence and hexahistidine tag allowed us to partially purify SLP2 from supernatant samples with immobilized metal ion affinity chromatography in sufficient quantities for experiments. In western blot analysis, a double band near the 37 kDa marker was seen in samples from the SLP2(12-469-his) expressing strain, but they were absent in similarly treated samples from the background strain (Fig. S5). This size is close to the size of the expressed fragment without the native I9 domain of SLP2 (35.9 kDa). A host cell protein, seen in SDS-PAGE and WB analysis at around the 75 kDa marker, co-purified with SLP2(12-469-his) and was also purified from the background strain samples (Fig. S5a) However, clearly higher protease activities were detected from the samples purified from SLP2(12-469-his) expressing strain than from the empty background strain (Fig. 2a). This protease activity was inhibited by the presence of a synthetic I9 inhibitor peptide (amino acids 82-151) and by serine protease inhibitors chymostatin or

**Fig. 2.**
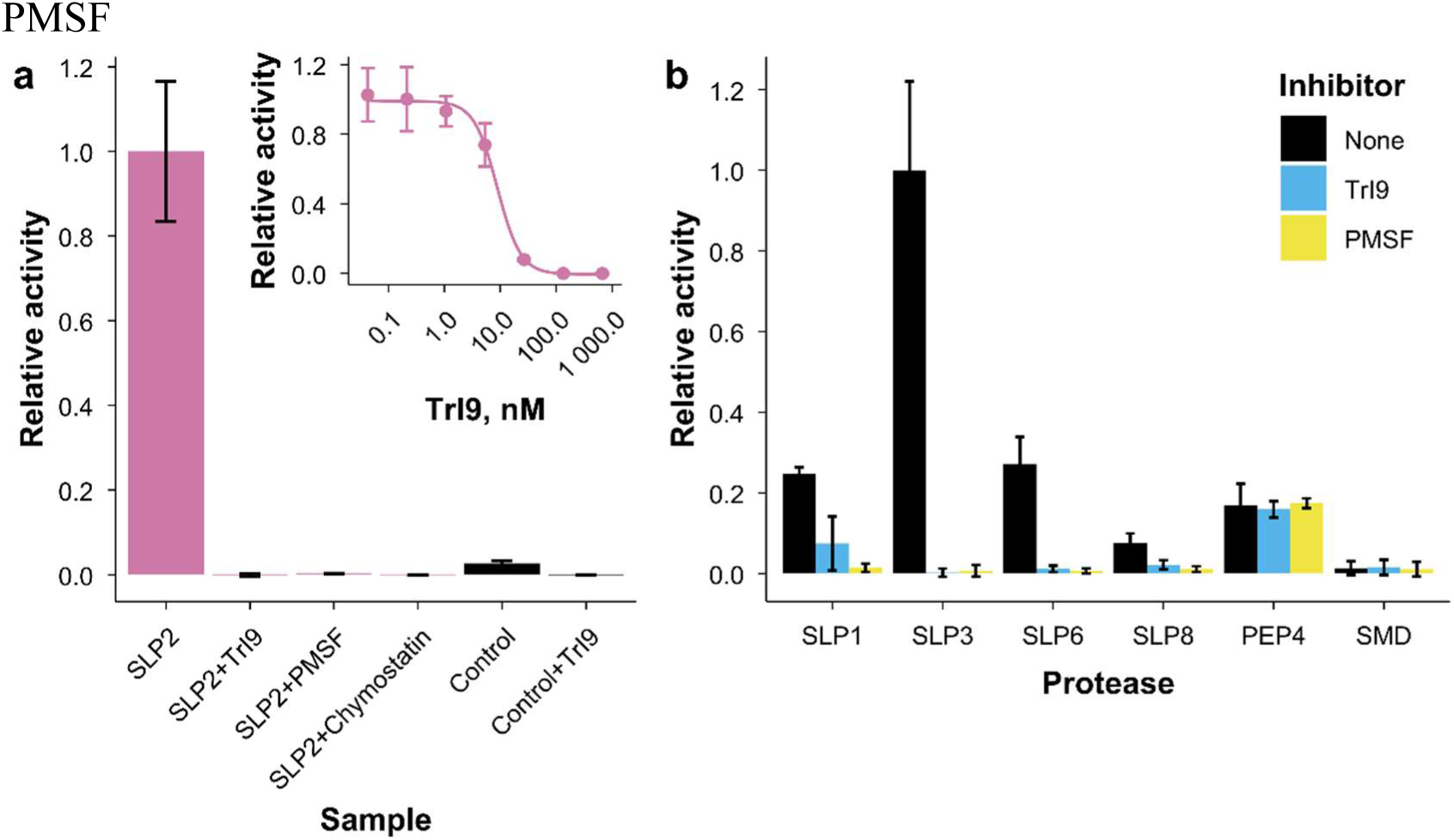
TrI9 can inhibit multiple *T. reesei* serine proteases. a) Protease activities of immobilized metal ion affinity chromatography purified *T. reesei* supernatant samples from SLP2(12-469_His)-expressing strain in the presence or absence of TrI9 or serine protease inhibitors PMSF or chymostatin. Similarly purified sample from the parental strain was included as control. The inset shows protease activities of partially purified SLP2(12-469_His) sample as a function of TrI9 concentration, with the line showing four parameter logistic curve fit. The TrI9 results shown in the main figure are with the highest concentration shown in the inset. Activities were normalized to trypsin control sample in each experiment and are shown relative to activity of the SLP2 sample. b) Protease activities of culture supernatants from *P. pastoris* strains expressing *T. reesei* SLP1, SLP3, SLP6, or SLP8 in the presence or absence of 100 nM synthetic TrI9 peptide. Supernatants from the parental strain (SMD) and a PEP4-expressing strain were included as controls. PMSF was used as control serine protease inhibitor. Activities are shown relative to activity of the SLP3 sample. Bars and points represent mean, and error bars standard deviation between two (a) or three (b) assays.

Based on computational modelling (Fig. S4), TrI9 is predicted to bind and potentially inhibit other SLPs in addition to SLP2. To test this experimentally, we produced recombinant SLP1, SLP3, SLP6 and SLP8 in *P. pastoris* and assessed if their protease activities could be inhibited with the synthetic TrI9 peptide (Fig. 2b). An empty background strain, a PEP4-producing strain, and a chemical protease inhibitor PMSF were used as controls. TrI9 appeared to inhibit the activity of all tested SLPs but had no effect on the activity of the aspartic protease PEP4. Weaker inhibition was observed for SLP1 and SLP8 at the tested TrI9 concentration. This is consistent with the structural modelling, in which SLP1 yielded the lowest confidence metrics among all tested SLPs (Fig. S4), suggesting it is the weakest target of TrI9 in the tested panel.

### Overexpression of TrI9 stabilizes CBM-AQ12-CBM against protein degradation *in vivo*

Deletion of *Slp2* improved the stability of CBM-AQ12-CBM; however, as expected, the deletion strain had growth and sporulation defects (Fig. S2). To study if TrI9 is effective in inhibiting the SLP2 activity also *in vivo* when co-expressed with CBM-AQ12-CBM and could improve its stability, different fungal strains were generated. To generate TrI9 overexpression (OE) strains, two versions of varying sequence length (with and without the intrinsically disordered residues 1-80) were expressed under the xylanase 2 (*xyn2*) promoter and targeted for secretion either by fusing the TrI9 with either the hydrophobin 1 (*hfb1*) signal sequence or XYN2 carrier protein. These genetic elements were chosen to ensure efficient secretion of the TrI9 protein to the culture medium. To study the possible biological role and effect of the inhibitor, the *trire_122168* gene was deleted from the *T. reesei* genome. Transformants were selected based on PCR screening for integration of the TrI9 cassette or deletion of the target *TrI9* gene, and the production levels of CBM-AQ12-CBM were analysed from twelve transformants from each construct using 24-well plate cultivations. The best performing strains were purified and chosen for future analysis (data not shown).

The best strains from the initial screen were grown in 24-well plates and samples were collected after four days of cultivation. The production levels and the stability of the CBM-AQ12-CBM protein were analysed by western blot utilizing the C-terminal Strep II tag in the CBM-AQ12-CBM (Fig. 3). The parental strain carrying the native *trire_122168* gene as well as the *TrI9* deletion strains showed only a weak signal of the full-length protein. All the TrI9 OE strains showed improved protein stability, and full-size CBM-AQ12-CBM target protein could be detected in the culture supernatants. Two different lengths of TrI9 sequence were tested. The short version (residues 82-151) was shown to be effective in *in vitro* tests. The full-length TrI9 (residues 1-151) did not show improved effect on CBM-AQ12-CBM stability, but the XYN2 carrier fusion yielded higher levels of the target protein than expression utilizing the HFB1 signal sequence in 24-well plate cultures (Fig. 3b). However, the fragmentation pattern was still visible, indicating that inhibition of SLP2 activity toward the CBM-AQ12-CBM target protein was not complete. Because CBM-AQ12-CBM has N-terminal His-tag efforts to detect the TrI9 using an anti-His antibody proved inconclusive. Thus, we confirmed the overexpression of *TrI9* gene using RT-qPCR from xyn2 carrier TrI9 short strain from bioreactor fermentation (Fig. S6).

**Fig. 3.**
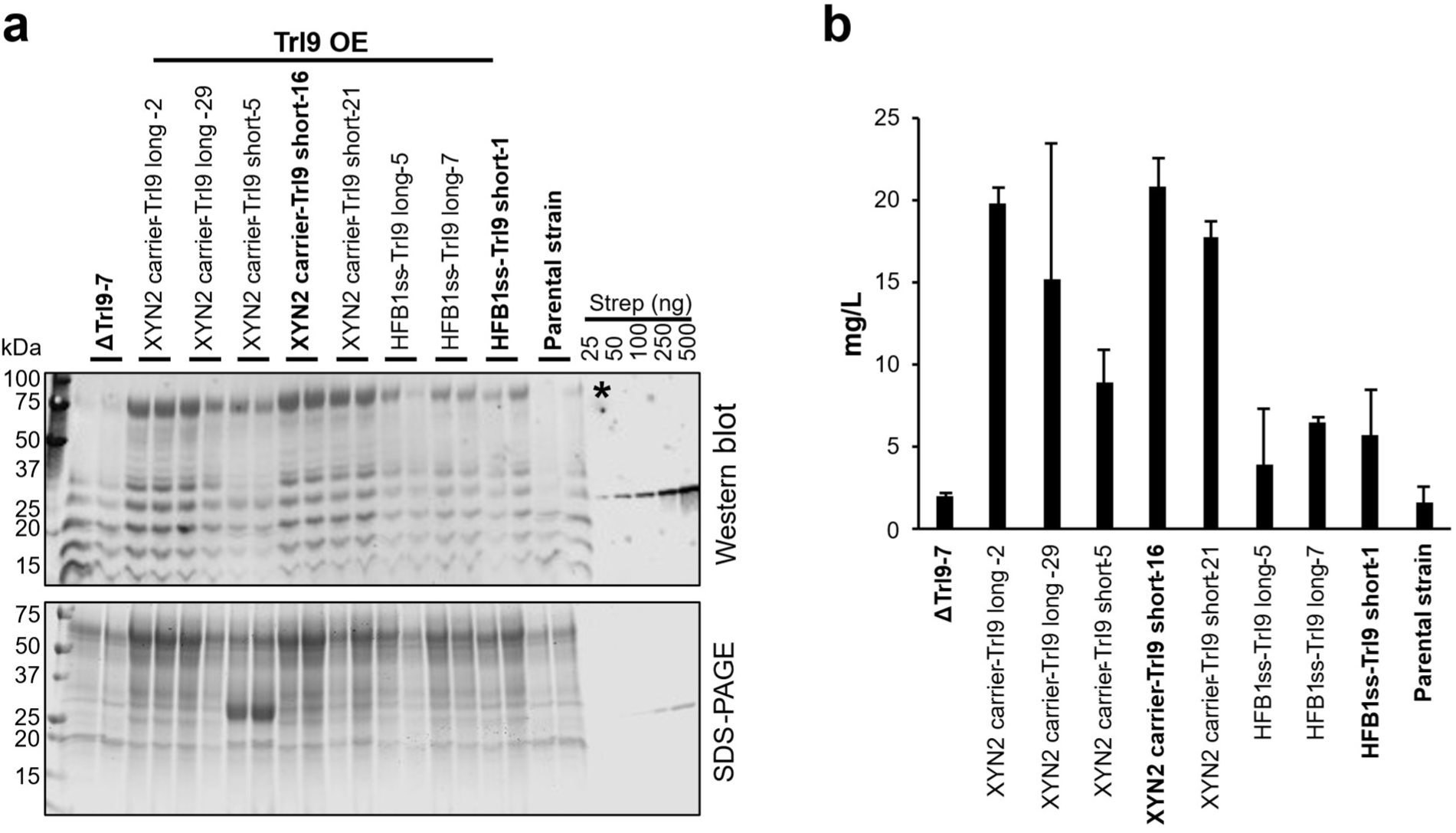
TrI9 overexpression stabilizes the protease-sensitive silk protein CBM-AQ12-CBM in small-scale cultures. *T. reesei* strains producing CBM-AQ12-CBM were cultivated in 24-well deep-well plates for 4 days and analyzed for target protein integrity. Overexpression of the subtilisin-like protease inhibitor TrI9 increased accumulation of full-length CBM-AQ12-CBM compared to the parental and ΔTrI9 strains. **a** Western blot detection of CBM-AQ12-CBM using the C-terminal Strep II tag (upper panel) with corresponding Coomassie-stained SDS-PAGE showing total secreted proteins (lower panel). Full-length CBM-AQ12-CBM (∼75 kDa) is indicated by an asterisk and lower-molecular-weight bands correspond to degradation products. **b** Quantification of intact CBM-AQ12-CBM based on western blot signal intensity using a GFP–Strep-tag II standard curve (25–500 ng). TrI9 was expressed using either XYN2 carrier or HFB1 signal sequence in long and short variants. Cultivations were performed in duplicate, and the experiment was repeated independently at least three times. Strains marked in bold was chosen for bioreactor cultivations.

### TrI9 inhibitor improves CBM-AQ12-CBM stability in bioreactor cultivations

The best performing TrI9 overexpression strains based on 24-well plate experiments were cultivated in Ambr® 250 High throughput bioreactors. As no difference was seen between the long and short versions of TrI9, we continued with the short TrI9 version with both XYN2-carrier and HFB1 signal sequence strains. All cultivations were carried out with the same batch media and same initial feed rate. The production of CBM-AQ12-CBM was analysed from bioreactor culture supernatants using SDS-PAGE and western blot, and the amount of full-length intact CBM-AQ12-CBM was quantified. No target protein was detected in the TrI9 deletion strain or the parental strain at pH 5 (Fig. 4). Both TrI9 OE and *slp2* deletion strains showed significant accumulation of the target protein. At pH 5 the TrI9 OE strains showed transient improvement of CBM-AQ12-CBM production which reached the peak at day 3-4 after which the levels started to decrease (Fig. 4b). The *slp2* deletion strain the peak production level was detected at day 5 and the inhibition was more stable than with TrI9 OE strains. In the bioreactors the HFB1 signal sequence construct of TrI9 performed better than the XYN2 carrier strain in contrast to 24-well plate cultures in which the opposite effect was observed. The highest titer of 0.5g/L was obtained with HFB1ss TrI9 strain, exceeding the amount observed in *slp2* deletion strain (Fig. 4b). The CBM-AQ12-CBM stability was further improved when pH was decreased from pH 5 to pH 4, resulting in 1.5 g/L of full length CBM-AQ12-CBM with XYN2 carrier (Fig. 4d). Furthermore, the inhibition was more stable over time.

**Fig. 4.**
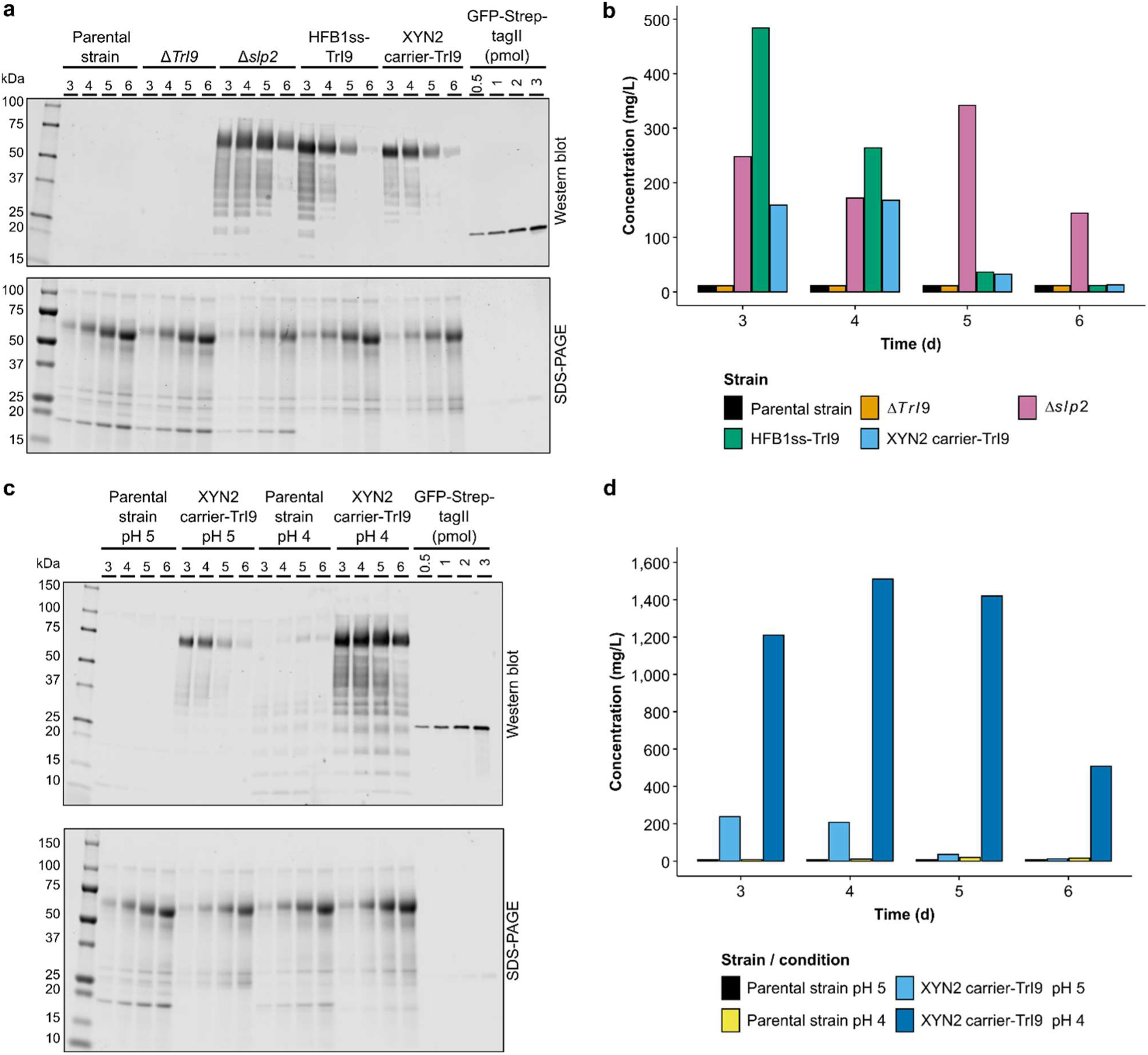
CBM-AQ12-CBM accumulation in TrI9 overexpression strains in bioreactor cultivations and a positive effect of lowering the cultivation pH. **a** Western blot analysis of culture supernatants from the parental strain, deletion strains (Δ*TrI9* and Δ*slp2*) and TrI9 overexpression strains (HFB1ss-TrI9 short and XYN2 carrier-TrI9) collected on days 3–6 (upper panel). Corresponding total protein profiles are shown by Coomassie stained SDS-PAGE (lower panel), 1 µl of culture supernatants were loaded. **b** Quantification results of intact CBM-AQ12-CBM in parental strain, deletion strains and TrI9 overexpression strains based on western blot (Fig. S7) from pH 5 cultivations. **c** Western blot analysis of culture supernatants from the parental strain and TrI9 overexpression strain (XYN2 carrier-TrI9) at pH 5 compared to the same strains at pH 4. Supernatants were collected on days 3–6 (upper panel). Corresponding total protein profiles are shown by Coomassie stained SDS-PAGE (lower panel), 1 µl of culture supernatants were loaded. **d** Quantification results of intact CBM-AQ12-CBM in parental strain and TrI9 overexpression strain at pH 4 and 5 based on western blot (Fig. S7). Western blot detection was performed using a primary antibody against Strep II tag of CBM-AQ12-CBM. A GFP-Strep-tag II control protein standard curve (0.5–3 pmol, corresponding to 30–180 ng of target protein) was included for quantification.

To validate the overall performance of selected stains in bioreactors, secretion of total protein levels were measured from culture supernatants (Table 2). At pH 5 the total protein secretion of the parental and TrI9 deletion strains was at similar level, thus resulting in similar bioprocesses. Lower cultivation pH had substantially higher target and total protein titer in contrast to the fermentation carried out at pH 5. In addition, the ratio of target and total proteins was higher at pH 4 implying that lower pH could lead into lower protein background in this case. Remarkable is, however, how cultivation at pH 4 resulted in a clearly higher biomass specific total protein yield in contrast to pH 5. This could be considered as an indication of a higher theoretical overall secretion capacity in contrast to pH 5 (Fig. 5).

**Fig. 5.**
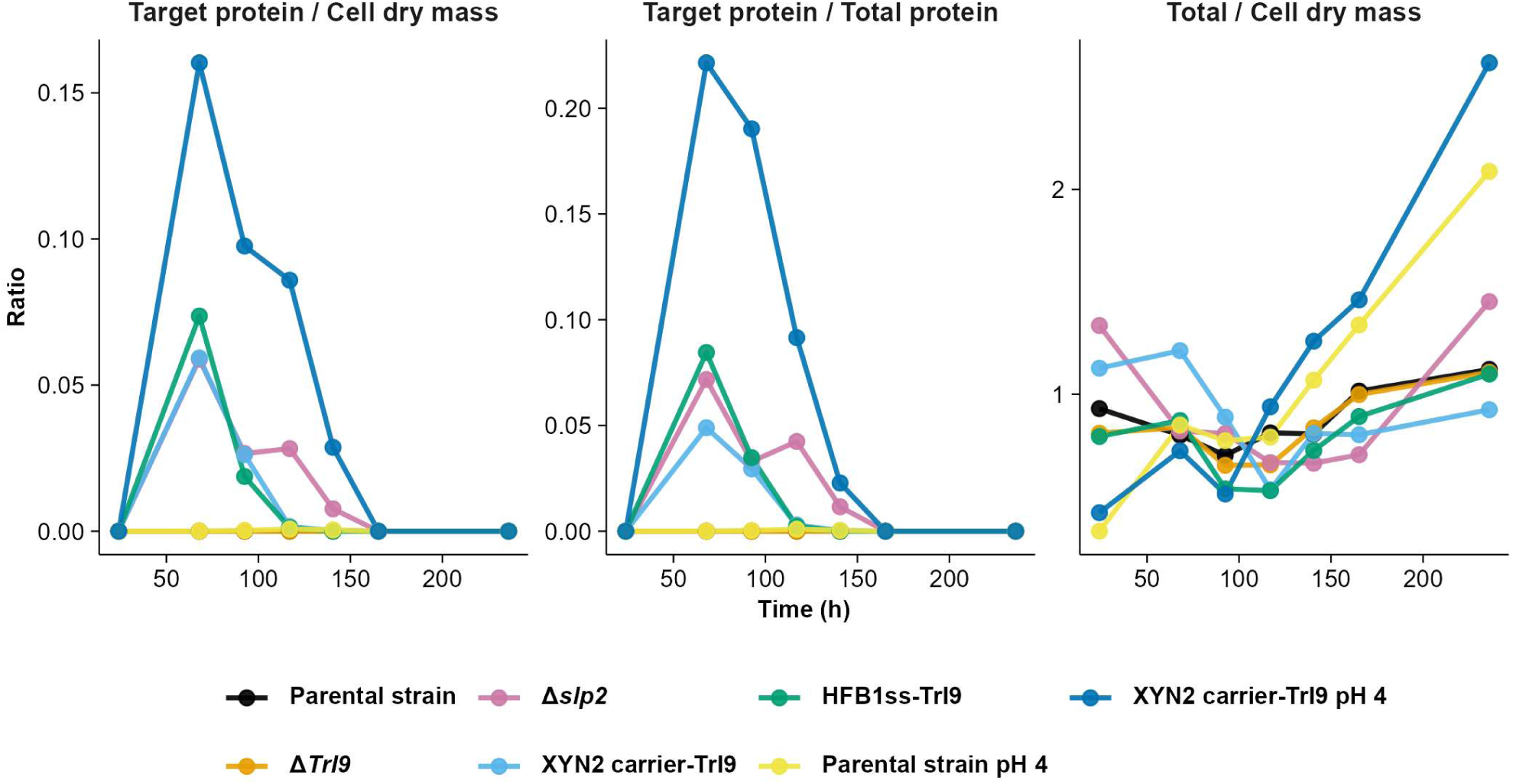
Evolution of background protein and biomass specific product amount. Decreasing the pH to 4.0 increased both biomass specific product formation and the quantity of product in relation to background proteins. Biomass was calculated by normalizing product formation to dry mass, where dry mass was estimated from reflectance measurements using robust linear regression. Target protein was under detection limit in *TrI9* deletion and parental strains at both pHs.

### TrI9 OE reduces the total protease activity in bioreactor cultivations

The total protease activity was measured from bioreactor culture supernatants (Fig. 6). The TrI9 OE strains had noticeably reduced total protease activity compared to both the parental and *TrI9* deletion strains. Only by deleting the *slp2*, protease activity was lowered approximately 75 % compared to the parental strain suggesting that it plays a substantial role in the residual protease activity within the 11x protease deletion background. The TrI9 OE strains had slightly lower total protease activity than *slp2* deletion strain indicating the ability to inhibit other serine proteases. Furthermore, total protease activity measured at pH 4 was lower than at pH 5 in both the parental strain and the TrI9 OE strain.

**Fig. 6.**
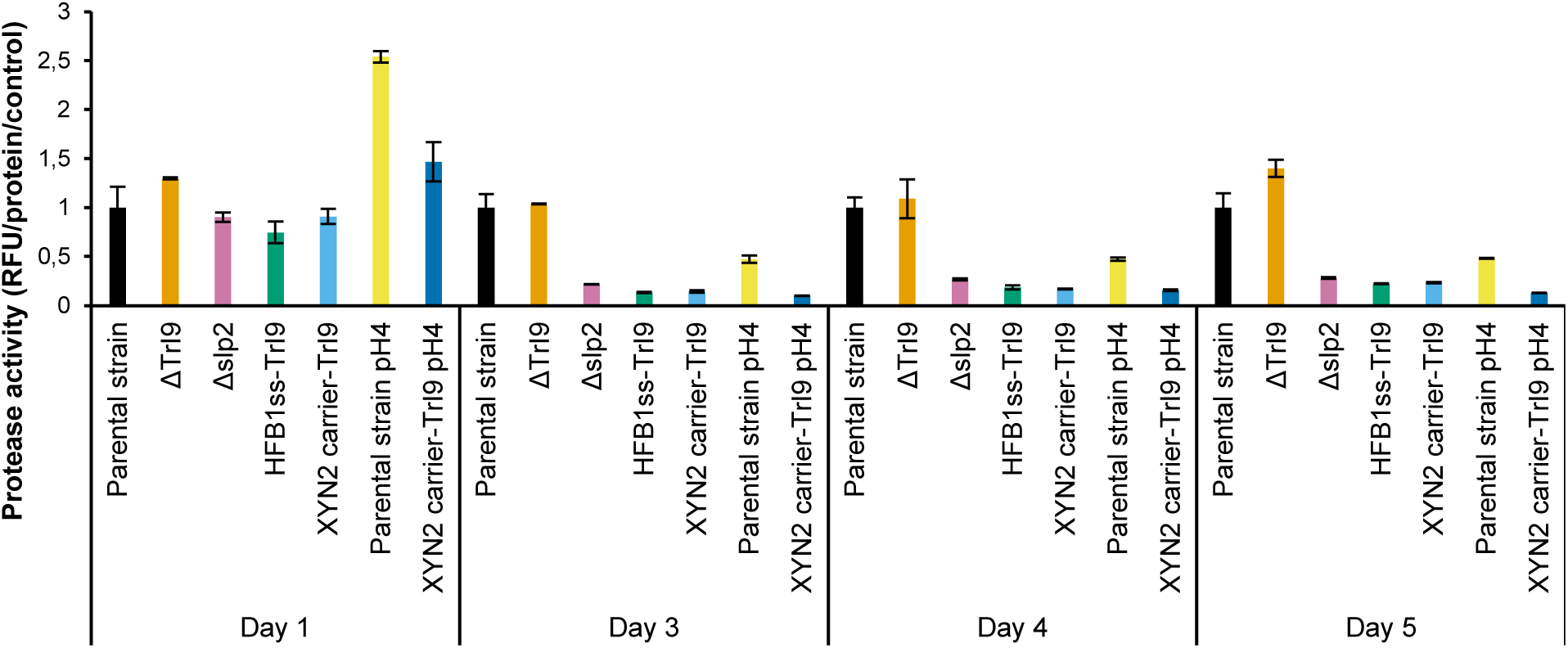
Overexpression of TrI9 reduces extracellular protease activity in bioreactor cultivations. Protease activity was measured in culture supernatants collected from bioreactor cultivations on days 1, 3, 4, and 5. Fluorescence after 60 min incubation with the fluorescent casein substrate was first normalized to the total protein concentration of each sample (Table S2) and then to the parental strain of each time point (set to 1). Unless otherwise noted, all strains were cultivated at pH 5. Strains shown: Parental strain, Δ*TrI9*, Δ*slp2*, HFB1ss-TrI9, XYN2 carrier-TrI9, Parental strain pH 4, and XYN2 carrier-TrI9 pH 4. Data represent the mean of three technical replicates from a single bioreactor cultivation.

### Overexpression of *TrI9* has no major effect on growth or sporulation of *T. reesei*

*Slp2* deletion had a negative effect on the growth and sporulation of *T. reesei*. Poor growth and sporulation not only limit strain performance and secretion levels but also cause challenges for future strain engineering efforts. Different methods were employed to investigate how the overexpression and deletion of *TrI9* influenced growth. Sporulation of TrI9 OE strains on PD plates was not visibly different from the parental strain (Fig. S8). Microscopic inspection revealed clear morphological differences as irregularly shaped spores of the *slp2* deletion strain, while the spores of the *TrI9* overexpression and deletion strains appeared round and similar to the spores of the parental strain (Fig. S9d).

To study spore germination and growth of *T. reesei* strains overexpressing the *TrI9* in more detail, their growth was monitored using automated time-lapse microscopy (oCelloScope) and compared to the parental strain and *slp2* and *TrI9* deletion strains (Fig. S9). Growth performance was quantified using multiple parameters: growth rate in the exponential phase, lag time duration, time to 50% germination and total growth (area under the curve, AUC) (Fig. S9a-c, Table S4). The *slp2* deletion strain showed consistently impaired performance across all quantified growth and germination metrics. Specifically, *slp2* deletion strain showed significantly reduced exponential growth rate, prolonged lag phase, slower germination and lower total growth (*p* < 0.01) (Fig. S9, Fig. S10).

In contrast, *TrI9* overexpression and deletion strains showed growth and germination behavior largely comparable to the parental strain. A statistically significant increase in the time to 50 % germination was observed for one *TrI9* overexpression strain, XYN2 carrier-TrI9 (Fig. S9b, Fig. S9c). In addition, slightly reduced exponential growth rate was observed for both *TrI9* overexpression strains (*p* < 0.05) and slightly shorter lag time for the *TrI9* deletion strain (*p* < 0.05); however, these observed differences did not meet the predefined significance threshold (*p* < 0.01). In particular, no significant differences in total growth (AUC) were detected for any of the TrI9 strains compared to the parental strain (Fig. S10). Full quantitative and statistical results are provided in Supplementary Tables S4–S6.

In bioreactor cultivations, deletion of the TrI9 or overexpression of HFBIss-TrI9 construct caused no changes in substrate consumption pace (lactose consumption), overall metabolism (CER) or in biomass build up (Reflectance) when compared to the parental strain (Fig. 7). However, XYN2 carrier TrI9 OE and the *slp2* deletion strains had clearly lower lactose consumption rate, growth speed, and biomass build up. It is noteworthy that despite the clear delay in overall metabolism, the gap in protein production and CER levels were diminished by the end of the fermentation. Based on a steadily decreasing CER observed at pH 4, a higher feed rate should be applied to improve the process at lower pH (Fig. 7).

**Fig. 7.**
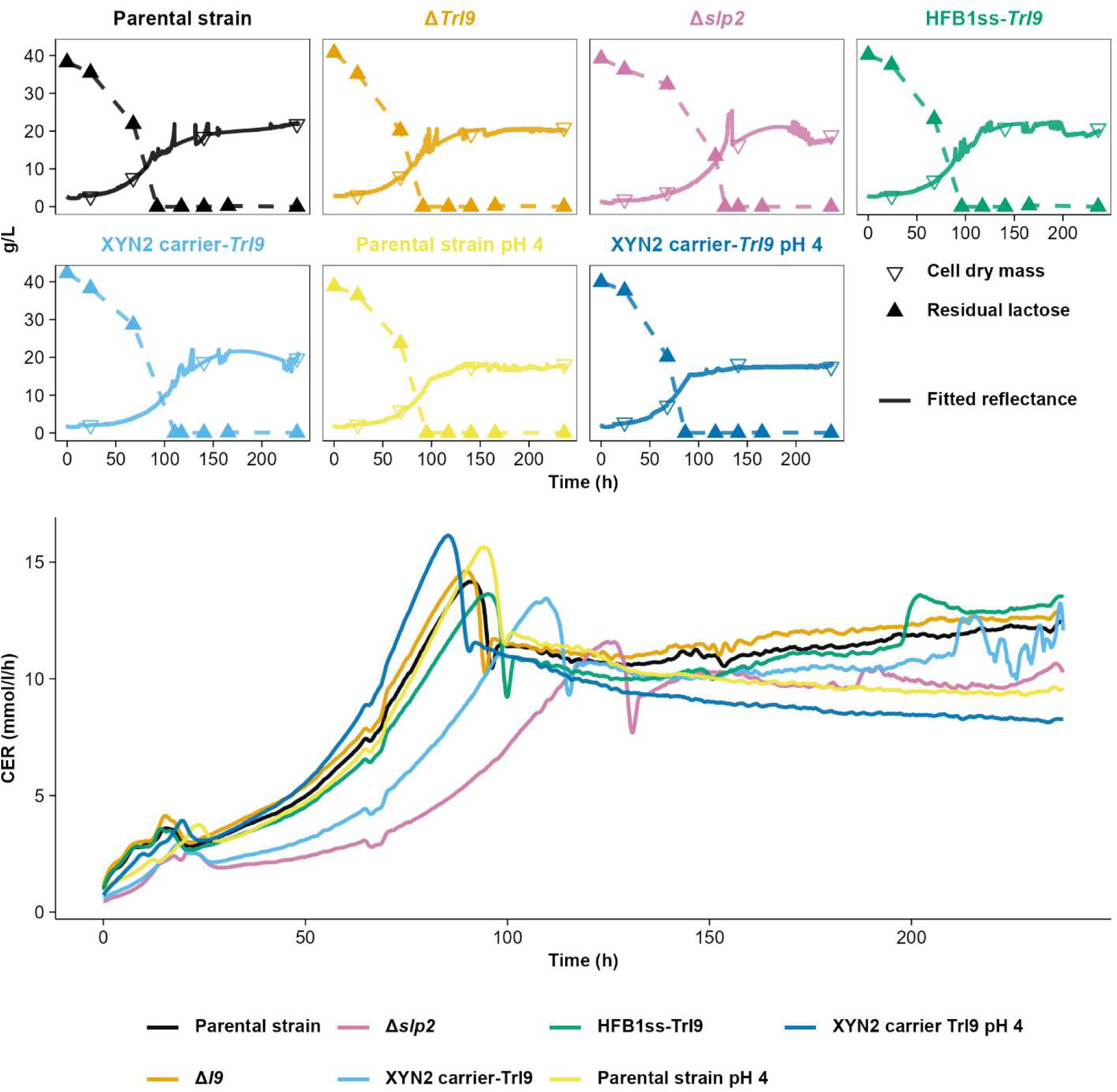
Substrate consumption, biomass build up and carbon evolution rate during bioreactor cultivation. Reflectance data was smoothed and interpolated using generalized additive models (GAMs, R). Reflectance signals were converted to dry mass using a robust linear calibration model fitted with the rlm function (MASS package, R). Residual lactose was measured by HPLC.

The parental strain and the XYN2-TrI9 OE were cultivated in pH 4 and pH 5. With both strains consumption of batch carbon was faster in pH 4 than in pH 5 (Fig. 7). Similar tendency was seen with XYN2-TrI9 OE in the pre-cultivation stage. Pre-cultivation at pH 5 produced only 1.6 g/l of biomass during the pre-cultivation, while the same strain cultivated at pH 4 yielded 2.3 g/l (Table 1).

## Discussion

We demonstrate here that protease activity in *T. reesei* can be substantially reduced by overexpressing and secreting inhibitory protein TrI9, encoded by *trire 122168* gene, and that this is a viable strategy to increase production yields of industrially relevant proteins. We demonstrated that the approach substantially reduced proteolysis of a highly sensitive silk protein CBM-AQ12-CBM both in small-scale cultures and in controlled bioreactor conditions and enabled maximum yields of 1.5 g/L. TrI9 protected also highly protease sensitive casein substrate in proteolytic assays, suggesting its applicability for protection of also other targets than silk proteins.

Importantly, TrI9 overexpression caused no major growth defects under most conditions; only the XYN2–TrI9 fusion strain showed growth reduction in a bioreactor cultivation at pH 5, which was not observed at pH 4. This contrasts with the severe phenotypes of *slp2* deletion strains, which exhibited reduced growth, delayed substrate consumption, and impaired protein secretion (this data and 5,10). This distinction underscores a practical advantage of inhibitor-based modulation over gene deletions: the native biological function of essential proteases is preserved, while extracellular degradation activity is selectively suppressed. This strategy provides a new biotechnological tool for stabilizing secreted protease sensitive recombinant proteins in culture supernatants, providing a solution for elimination of protease activities in filamentous fungi, a longstanding challenge in fungal biotechnology, without using costly commercial inhibitors or compromising strain fitness.

Computational structural analyses provide strong support for the functional role of TrI9 and offer mechanistic insight into its inhibitory activity. Structure-based search with FoldSeek identified TrI9 as a Pbi2 homologue, whereas sequence-based homology searches failed, highlighting the benefits of structure-based approaches in identifying functionally conserved proteins where sequence divergence has obscured evolutionary relationships. The predicted binding mode of TrI9 closely mimics that of the native SLP2 propeptide, with its C-terminal loop occupying the substrate-binding cleft and the P1 residue positioned in the catalytic pocket, placing TrI9 in the class of pure competitive inhibitors without prime-side contacts. In subtilisin-like proteases, the I9 propeptide serves a dual role as both an intramolecular folding chaperone and a latency-maintaining inhibitor that keeps the zymogen inactive during vacuolar trafficking, with release of the active protease triggered only by the acidic vacuolar environment (22, 23, 27, 28). The moderate energetic advantage of the native autoinhibitory domain over TrI9 observed in Rosetta calculations is therefore biologically logical: the propeptide must bind tightly enough to maintain protease latency throughout the secretory pathway, and the somewhat weaker affinity of TrI9 reflects its origin as a diverged paralogue rather than a co-evolved propeptide. This difference also suggests that affinity improvement through rational engineering of the TrI9 could be a productive direction for future work.

The computational analysis predicted that TrI9 can bind not only to SLP2, but also to all other eight annotated *T. reesei* SLPs, suggesting a more general inhibitory potential. This was confirmed experimentally for SLP1, SLP3, SLP6 and SLP8. Relative inhibitory strengths are also consistent with the computational predictions. SLP1 with the lowest confidence in the predicted TrI9–SLP1 binding geometry (ipSAE_min score (0.016), PAE (22.19 Å)) showed also the weakest inhibition experimentally compared to SLP3 (ipSAE_min score (0.762), PAE (4.59 Å)) and SLP6 (ipSAE_min score (0.739), PAE (4.96 Å)) with highest computational confidences. Although the number of tested proteases is too small to draw firm conclusions, this partial correspondence between the computational ranking and experimental inhibition strength is notable given that ipSAE_min has recently been identified as the best-performing individual metric for predicting protein–protein binding across a large and diverse benchmark dataset of experimentally validated binders (26), suggesting that its discriminative power may extend beyond de novo binder design to inhibitor–protease interactions of the type studied here. Although the Rosetta interface ΔG estimate for SLP1 appears comparable to the other SLPs, this value should be treated with caution since it is derived from a low-confidence structural model and is unlikely to reliably capture the true binding energy. The fragment docking analysis additionally identified a single high-confidence SLP2 substrate motif, PGQQ↓GPGQ, occurring up to 24 times across the full-length AQ12 domain, which is qualitatively consistent with the extreme proteolytic sensitivity of CBM-AQ12-CBM observed experimentally. It should be noted that the application of peptide-protein fragment docking to protease substrate identification has not yet been systematically validated in the literature, and the predicted cleavage sites should be regarded as suggestive rather than definitive as the actual cleavage site was not experimentally confirmed. In conclusion, structural modelling and experimental evidence suggest that TrI9 inhibits also other proteases than SLP2.

pH had a strong effect on the accumulation of CBM-AQ12-CBM and TrI9 inhibition. In bioreactor cultivations TrI9 overexpression strains accumulated full-length CBM-AQ12-CBM, whereas neither the TrI9 deletion nor the parental strain produced measurable intact product at pH 5. Lowering the cultivation pH to 4 further improved stability, consistent with previous reports that acidic conditions reduce serine protease activity in *T. reesei* (own data and (29, 30)). Based on the data, we hypothesize that at pH 5, TrI9 must compete with a more active and potentially more abundant subtilase pool, consistent with the residual fragmentation observed in 24-well plate and bioreactor experiments. Other plausible explanations are that the pH influences the binding affinity of the TrI9 to the proteases or that the TrI9 or the protease gene expression or protein abundance is affected by the pH. It was seen that especially at pH 5 the inhibition was transient and started to decrease after 3-4 days of cultivation in bioreactors. Similar phenomenon has been reported in yeast, where it was shown that subtilisin degraded the YIB2 inhibitor over time (31).

TrI9 overexpression and secretion increased accumulation of full-length CBM-AQ12-CBM in 24-well plate cultures and bioreactors; although degradation fragments were still detectable, intact protein was consistently present across all TrI9 overexpression strains, indicating partial but meaningful protection from proteolysis. A clear direction for further improvement is to enhance the secretion efficiency of the TrI9 and the stability of the inhibitor itself and its interaction with the SLP2 protease. These approaches will increase the effective concentration of TrI9 in the culture supernatant and strengthen the protection against SLP2-mediated degradation. Future work will therefore focus on evaluating additional native or engineered secretion signals and carrier proteins to identify variants that promote higher extracellular accumulation of the TrI9 without compromising folding or stability. To optimize expression levels placing the TrI9 under an inducible or condition responsive promoter would allow temporal expression control so that production would be activated primarily during the target protein production phase. Such inducible systems would reduce metabolic burden during early growth, improve strain robustness, and minimize potential fitness costs while still enabling strong inhibitory action at the stage where protease activity is most detrimental.

Additional strategy for inhibition improvement involves increasing the stability of the TrI9 – SLP2 interaction in the culture supernatant. Although the endogenous inhibitor is structurally well-suited to block SLP2 type proteases, its target affinity and resistance to extracellular degradation could likely be strengthened. In *P. ostreatus*, it was shown that different mutations in PoIA1 can improve or reduce its protease inhibition or chaperone activities (32). Hence, rational engineering of TrI9 by stabilizing key C-terminal residues that occupy the SLP2 catalytic cleft, may improve binding efficiency under process conditions. Additionally, fusing the inhibitor to stabilizing domains, modifying surface residues to resist proteolysis, or engineering variants with improved pH tolerance could further enhance *in situ* inhibitory capacity. Such approaches may be especially beneficial at higher pH, where SLP2 activity is maximal and proteolytic degradation of target proteins is augmented. Altogether, these strategies offer complementary routes toward creating a more robust TrI9 based inhibition system capable of sustained target protein protection in industrial fermentations.

Although we do not know the exact biological function of the TrI9, the presence of an independently encoded I9 inhibitor in the *T. reesei* genome suggests an additional level of regulation of the activity of subtilisin-like protease, beyond the intramolecular I9 propeptides that control folding and zymogen latency. Standalone I9 inhibitors in fungi (e.g., yeast PBI2 and *P. ostreatus* PoIA proteins) have been linked to protease regulation and, in some cases, broader vacuolar functions by binding the vacuolar SNARE complex and stabilizing the Vam3p SNARE protein (16, 31, 33-35). In yeast and *P. ostreatus* inhibitors were shown to function as chaperones facilitating proper folding of their cognate proteases (16, 31). Defining TrI9 expression, localisation and interaction partners, and testing Δ*TrI9* under nutrient limitation or other protease-inducing regimes, will help clarify its biological function.

Our results demonstrate the power of the integration of advanced computational technologies with biotechnology. The practical advance of structure-guided docking and protein-protein interaction analyses tools not only validate experimental findings but also accelerate experimentation which has traditionally been based on trial-and-error processes, facilitate the design of novel synthetic molecules, and possibly enable in the future the development of new predictive tools for protease susceptibility. Taken together, these findings highlight the potential of the TrI9 inhibitor as a versatile strategy to enhance the stability of recombinant proteins in *T. reesei* production system. In contrast to protease gene deletions which can yield strains with poor growth, impaired secretion, or unfavourable metabolic profiles, the expression of the TrI9 inhibitor can offer a finetuned means of reducing proteolytic activity without compromising host viability. The broad-spectrum inhibitory potential also raises intriguing possibilities for engineering strains with improved performance across diverse recombinant targets. Finally, the discovery of structurally related inhibitor genes in other fungal species, including the previously described *Aspergillus niger* PepC inhibitor (WO2009071530) (36), suggests that inhibitory proteins may be widely conserved and could represent an underexplored class of tools for fungal strain engineering.

## Materials and Methods

### Identification of TrI9 by structure-based homology search

To identify a putative I9-type inhibitor in the *T. reesei* genome, a structure-based homology search was performed using FoldSeek (20). The AlphaFold-predicted structure of *S. cerevisiae* Pbi2 (UniProt accession P0CT04; AlphaFold Protein Structure Database entry AF-P0CT04-F1-v6) was used as the query in a search against the AlphaFold/UniProt50 v6 database using the 3Di/AA combined scoring mode with a taxonomic filter restricted to *Trichoderma reesei*. TrI9 (encoded by *trire2_122168*; AlphaFold entry AF-G0RKY8-F1-model_v6) was returned as a hit (search probability 1.0, score 207, E-value 3.06 × 10⁻⁴), confirming structural homology despite limited sequence identity.

### AlphaFold3 structure prediction of inhibitor–protease complexes

All protein–protein and protein–peptide complex structures were predicted using a local installation of AlphaFold3 v3.0.1 (21) with the AlphaFold-beta-20231127 model parameters. Predictions were run with num_seeds = 5, num_diffusion_samples = 5, and num_recycles = 10. Multiple sequence alignments (MSAs) were generated prior to prediction using a local installation of ColabFold v1.5.5 (37) with MMseqs2 v18.8cc5c (37) against the combined ColabFold MSA database comprising UniRef30 and environmental (metagenomic) sequences. The resulting a3m MSA files were provided directly as input to AlphaFold3. For each complex, the highest-confidence predicted structure (ranked by AF3 confidence metrics) was selected for downstream analysis.

Predicted complexes included: full-length SLP2 (used to characterise the endogenous autoinhibitory I9 domain configuration); TrI9 co-folded with the SLP2 catalytic domain alone (residues 153–474); and TrI9 co-folded with the isolated catalytic domains of each of the remaining eight annotated subtilisin-like proteases (SLPs) encoded in the *T. reesei* QM6a genome (SLP1–SLP10; gene identifiers following the trire2 annotation).

### Interface confidence metrics

Interface confidence was assessed using two complementary metrics extracted from AlphaFold3 predictions. The interaction prediction Score from Aligned Errors (ipSAE_min) was computed following Overath et al. (26) and Dunbrack (38). ipSAE is calculated analogously to ipTM but restricts the calculation to interchain residue pairs with predicted aligned error (PAE) below a cutoff of 10 Å, and dynamically scales the d₀ parameter with the square root of the number of residues within this cutoff, thereby penalising small interfaces. The asymmetric ipSAE score was computed in both directions (inhibitor → protease and protease → inhibitor), and the minimum of these two values (ipSAE_min) was taken as the final metric, as it captures the weakest-link interaction direction and has been shown to be the strongest individual predictor of in vitro binding across diverse protein–protein interaction datasets (26) The mean interface PAE was computed as the average of all interchain predicted aligned error values between the inhibitor and protease chains, providing an overall measure of confidence in the predicted binding geometry. Both metrics were calculated using the analysis pipeline of Overath et al. (26), available at https://github.com/DigBioLab/de_novo_binder_scoring.

### Rosetta-based interface energy calculations

Interface binding energies were estimated using PyRosetta (39) following the protocol implemented in Overath et al. (26), which was adapted from the BindCraft scoring pipeline (40). Briefly, each AF3-predicted complex structure was first subjected to constrained structural relaxation using the FastRelax protocol (41) with the ref2015 energy function (24). Interface binding free energies (ΔG, in Rosetta Energy Units, REU) were then estimated using Rosetta’s InterfaceAnalyzerMover, which calculates the energetic cost of separating the two chains after repacking interface residues in the unbound state. The Rosetta ΔG values thus reflect the predicted energetic stability of the complex interface and are used here for comparative ranking of TrI9 binding to different SLP targets. All computed metrics are reported in Fig. S2.

### Computational prediction of SLP2 cleavage sites in AQ12 silk domain

To identify putative cleavage sites of the *T. reesei* SLP2 protease within the AQ12 artificial spider silk domain, an AlphaFold2 multimer-based fragment docking approach was employed, following the fragmentation strategy described by Lee et al. (2024) (18). The AQ12 domain consists of twelve tandem repeats of the sequence AAAAAAGGYGPGSGQQGPGQQGPGQQGPGQQGPGQQGPYGPGAS. A two-repeat unit (AQ2) was used as a representative query sequence, as the periodicity of the repeat renders it sufficient to sample all sequence contexts present in the full-length domain.

The AQ12 sequence was fragmented into 10-residue peptides using a sliding window with a frame shift of 3 residues, generating a set of overlapping fragments that collectively provide dense coverage of the entire repeat sequence. Each fragment was submitted for complex structure prediction with the catalytic domain of SLP2 (*trire2_123244*) using AlphaFold2 Multimer v3. Multiple sequence alignments (MSAs) for the input sequences were generated using the same pipeline as described for AlphaFold3 predictions in the preceding methods section. Five structural models were generated per fragment-protease pair.

The confidence of each predicted fragment-protease complex was assessed using the actifpTM metric (19), a modified version of the standard ipTM score that corrects for bias introduced by flexible flanking regions by restricting the pTM calculation to residues participating in the predicted interface. This correction is particularly important for short peptide substrates, where flanking sequence context can substantially deflate the standard ipTM value even when the core binding interaction is accurately predicted. A threshold of actifpTM > 0.85 was applied to identify high-confidence docking models, following the calibration established for peptide-protein interactions by Varga et al. (2025) (19).

Models passing the actifpTM threshold were subsequently subjected to structural inspection to evaluate whether the bound fragment configuration was compatible with catalytic cleavage, specifically assessing positioning of the putative scissile bond relative to the catalytic triad of the SLP2 catalytic domain. Fragments meeting both the confidence and geometric criteria were used to define putative cleavage motifs within the AQ12 repeat sequence.

### Protein structure visualisation

All protein structures were visualised and figures were prepared using UCSF ChimeraX v1.10.1 (42).

### Construction of *T. reesei* strains used in the study

*Escherichia coli* DH5α (Invitrogen, Thermo Fisher Scientific, USA) was used for propagation of the plasmids. *Trichoderma reesei* strain that has 11 secreted proteases deleted (43) was further modified by transforming a spider silk AQ12 expression cassette from pAWP188 (43). The pAWP188 vector contained cbh1 promoter and terminator and hygromycine marker with CBHI carrier fused to CBM-AQ12-CBM fragment. AQ12 contains 12 repeats derived from ADF3, which is one of the major dragline silk proteins produced by European garden spider (*Araneus diadematus*) (44). The flanking carbohydrate binding motives (CBMs) were derived from cellobiohydrolases 1 and 2 (Fig. S11) and obtained strain was used in overexpression of the TrI9 cassettes.

The overexpression vectors were ordered from Genscript, China. The expression vector pSUFU131 contained *Xyn2 carrier-TrI9* long gene fragment with his-tag, pSUFU132 contained *Xyn2 carrier-TrI9* short gene fragment with his-tag, pSUFU133 contained Tr*I9* long gene with *hfb2* signal sequence and his-tag, and pSUFU134 contained *I9* short gene with *hfb2* signal sequence and his-tag. All vectors contained also *pyr4* marker and sequences targeting the expression constructs to *xyn2* locus in *T. reesei* genome. The expression cassettes were liberated from the plasmid with MssI restriction enzyme and purified from an agarose gel and transformed into *T. reesei* with protoplast method essentially as described in (45).

TrI9 and Slp2 deletion strains were generated with CRISPR-Cas9 as described in (46) using pair of gRNAs targeted to 5’ and 3’ ends of the gene. Guide RNAs used in the deletions are listed in Table S3. The integration of the expression cassettes or the deletion of the genes was confirmed using PCR and screened in 24-well plates. The best performing strains single spore purified.

SLP2 expression vector contained SLP2 with the predicted native signal sequence (amino acids 1–15) replaced by CBH1 signal sequence (amino acids 1-17 from trire2_123989) and with the C-terminal domain (470–534) replaced by hexahistidine tag. SLP2 was expressed with its endogenous promoter and the construct contained flanks for integration into the endogenous locus as well as Pyr4 selection marker. The SLP2 expression construct was transformed into a Pyr4^-^ background strain with 3 additional protease deletions to the previously mentioned 11 protease deletion strain (gap2/trire2_106661, pep9/trire2_79807 and amp5/trire2_122083). Pyr4^+^ parent of the 14 protease deletion background strain was used as a control in SLP2 purification experiments.

The remaining *T. reesei* proteases, SLP1, SLP3, SLP6, SLP8 and PEP4 were produced in *P. pastoris.* The sequences were ordered codon optimized and cloned in pPICZαA vector from Genscript, China, and produced as C-terminally His-tagged under the methanol-inducible AOX promoter with α-factor secretion signal peptide in *Pichia pastoris* SMD1168H (Invitrogen, Thermo Fisher Scientific, USA). Proteins were produced under MeOH induction as described in (47) for two days and cells were removed by centrifugation and supernatant stored at -20 °C.

### RNA extraction and RT-qPCR

The attempts to detect TrI9 with western blot using His-tag were inconclusive. The small size of TrI9 made the detection difficult and several contaminating bands from His-tagged CBM-AQ12-CBM were detected and the origin of the signal was uncertain (data not shown). Thus, RT-qPCR was done to verify the overexpression of the TrI9.

Mycelium samples for RT-qPCR analysis were collected from Ambr^®^ bioreactor cultures by vacuum filtration and frozen immediately with liquid nitrogen. RNA was extracted from the samples with RNeasy plant mini kit (Qiagen, Hilden, Germany) following the manufacturer’s protocol for purification of total RNA from plant cells and tissues and filamentous fungi, treated with DNAse I (Thermo Fisher Scientific, Waltham, MA, USA) and reverse transcribed into cDNA (Transcriptor First Strand cDNA Synthesis Kit, Roche, Basel, Switzerland). RT-qPCR was performed with LightCycler^®^ 480 SYBR Green I master mix (Roche) with CFX Opus Real-Time PCR system (Bio-Rad, Hercules, CA, USA), with two technical replicates per sample. The used primers are listed in Table S3, and their amplification efficiencies were determined by measuring C_q_ values from a serially diluted cDNA sample. Expression levels were calculated with the 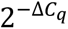 method (48) by correcting for amplification efficiency, using equation 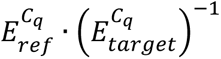 (where *E* = efficiency, *C*_*q*_ = quantification cycle and 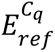 = geometric mean of reference gene 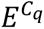 values), with *sar1* and *act* as reference genes (49).

### CBM-AQ12-CBM protease-sensitivity assay using *T. reesei* supernatants and protease inhibitors

The following stock components were used for the assay: lyophilized CBM-AQ12-CBM with a Strep-tag (1 mg/ml in 50 mM sodium citrate), assay buffer (50 mM Tris-HCl pH 4.8), *T. reesei* supernatant from 11 protease deletion strain Ambr cultivation with 4 % glucose and 1 % yeast extract, pH4.8, 5 days cultivation, 20 mg/ml total protein concentration), Dimethyl sulfoxide (DMSO), Phenylmethylsulfonyl fluoride (PMSF) 100 mM in DMSO, Iodoacetamide (IAA) 100 mM in MQ, Ethylenediaminetetraacetic acid (EDTA) 0.5 M in water, Chymostatin (CHY) 4 mM in DMSO, Pepstatin A (PepA) 4 mM in DMSO, Soybean trypsin inhibitor (SBTI) 40 mg/ml in water. The assay mix contained 83 µl assay buffer, 6 µl CBM-AQ12-CBM, 10 µl *T. reesei* culture supernatant and inhibitor additions of 0.5 µl (PepA, SBTI), 1 µl (CHY, EDTA, PMSF) or 2 µl (IAA). As controls, supernatant was replaced by assay buffer, and inhibitors were replaced by 2 µl DMSO or water. Mixtures were incubated at 28°C, and samples for SDS-PAGE were taken after 1.5h, 3.5h and 21h after initiation of the treatment. CBM-AQ12-CBM was detected by western blotting using Recombinant Rabbit Monoclonal Antibody (Cell Signaling Technology, MA, USA) followed by an IRDye 800CW anti-rabbit secondary antibody (LI-COR Biosciences, Nebraska, USA).

### Strain cultivations in 24 deep well plates

*Trichoderma reesei* strains were cultured on potato dextrose agar (PDA) plates at 25 °C for 7 days to allow for sporulation. Spores were harvested in (0.8 % NaCl, 0.025 % Tween-20, 20 % glycerol) and quantified using a Luna-II Automated Cell Counter (Logos Biosystems, South Korea). Spore suspensions were diluted to a final concentration of 1 × 10⁷ spores/mL in 3.5 mL of growth medium (4 % lactose, 2 % spent grain extract, 15g/l KH_2_PO_4_, 5.4 g/l Na_2_SO_4_, trace elements, 100 mM PIPPS di-ammonium citrate, 2.4 mM MgSO₄, 4.1 mM CaCl₂, pH 4.8) and were transferred into individual wells of a 24-well deep-well plate and incubated in a Multitron shaker (Infors HT) at 28 °C and 800 rpm for 4 days. After incubation, 250 µL of culture supernatant from each well was transferred to a filter plate (#MSDVN6550, Millipore) and centrifuged at 1,800 × g for 8 min to remove cells prior to downstream analyses. Microscope images were taken with Leica DM 2000 microscope (Leica Microsystems, Wetzlar, Germany).

### SDS-PAGE, western blot and protein quantification

Protein samples were resolved by SDS–PAGE using 4–20 % Criterion™ TGX™ Precast Midi Protein Gels (Bio-Rad, Hercules, CA, USA) together with Prestained Protein Ladders (Bio-Rad, Hercules, CA, USA). For Coomassie-stained gels, visualization was performed using PageBlue Protein Staining Solution (Thermo Scientific, USA) according to the manufacturer’s instructions. For western blotting, proteins were transferred to nitrocellulose membranes using the Trans-Blot® Turbo™ Transfer System (Bio-Rad, Hercules, CA, USA). To detect CBM-AQ12-CBM α-Strep primary antibody (IBA Lifesciences, Göttingen, Germany) 1:1000 and anti-mouse secondary antibody 1:30 000 (Li-COR Biosciences, Nebraska, USA) were used. Commercial GFP-Strep-tagII control protein was used as standard for protein quantification from the western blots (IBA Lifesciences, Göttingen, Germany) Coomassie-stained gels and western blots were imaged and quantified by using an Odyssey CLx Imaging System (Li-COR Biosciences).

### Pre cultivation and bioreactor cultures

Pre-culture was initiated with 1x10^7 spores /50ml and cultivated in preculture media containing 40 g/l lactose, 2 % spent grain extract, 15 g/l KH_2_PO_4_, 5 g/l(NH_4_)_2_SO_4_, 2.4 mM MgSO₄, 4.1 mM CaCl₂ and 1ml/l trace element solution (5 g/L FeSO4⋅7H2O, 1.6 g/L MnSO4⋅H2O, 1.4 g/L ZnSO4⋅7H2O, 3.7 g/L CoCl2⋅6H2O). Pre-cultures were cultivated for 48 h at 28 °C and 200 rpm on an orbital shaker (orbit diameter 25 mm), after which 1/3 volume of inoculum was added to fresh media and grown for additional 48h. Each strain was concentrated by centrifuging the inoculum at 4000 g for 10 min. Supernatant was removed and 18 ml of fresh process media was added to resuspend the pellet. The resuspended inoculum was used fully to gain 10 % inoculum volume.

Bioreactor run was carried out in Ambr® 250 High Throughput-system (Sartorius, Germany) system. The batch media consisted of 15 g/l of KH2PO4, 12 g/l (NH4)2SO4, 333 ml/l of a 6 % Spent grain extract, 0.6 g/l of MgSO4, 0.6 g of CaCl2, 1 ml/l of J673A antifoam, 1 ml/l of trace element stock solution and 40 g/l of lactose as the main carbon source. Trace element solution consisted of 10 g/l of FeSO4, 3 g/l of MnSO4, 3 g/l of ZnSO4, 4 g/l of CoCl2 1 ml/l of Verduyn vitamin solution. Citric acid was used to dissolve the metal ions. The cultivation was carried as a fed-batch fermentation where 20 % lactose feed was initiated by an automated system after a 10 % drop in CO2 % was detected. Based on the data the feed was initiated 2.9-4.05 h after the initial indication of lactose limitation. All reactors were supplemented with additional nutrients from day 5 onwards. Each day 0.2 ml of trace element stock, CaCl2 stock of 2.5 M and (NH4)2SO4 stock 500g/l were added to each reactor.

HPLC analyses were carried out as described in Jämsä et al. (50). Dry weight analyses were carried out by measuring either 5 or 10 ml of broth to a pre weighed Whatman gf/f filter paper. Broth was dried in 105 °C for 48 hours.

### Total protein measurement and protease activity assay

Protein concentration was quantified using the DC Protein Assay Kit (Bio-Rad, Hercules, CA, USA) according to the manufacturer’s instructions using BSA as a standard.

Protease activity in culture supernatants was quantified using the SensoLyte Red Protease Assay Kit (AnaSpec, Fremont, CA, USA). The fluorescently labelled casein substrate was diluted in assay buffer and used in the recommended ratio of 1 µl of substrate per 100 µl reaction. Unless otherwise noted, the 2x buffer provided by the kit was used as the assay buffer. The increase of fluorescence (Ex 546 nm, Em 575 nm) from the cleaved casein was kinetically monitored using Varioskan Flash plate reader (Thermo Fisher Scientific, USA) for at least 1 hour. Activities were determined as the slope of the linear portion of the time vs fluorescence curve, unless otherwise specified. Samples with assay buffer alone were used as negative control.

### oCellscope cultivation and data analysis

Fungal growth phenotype and spore germination were analyzed using automated time-lapse microscopy (oCelloScope, BioSense Solutions ApS, Farum, Denmark). Strains were grown on PDA plates at 28 °C until sporulation occurred and harvested and quantified as described above for the 24-well plate cultivation experiments. Spore suspensions were frozen and stored at -80 °C until use.

Fungal strains were cultivated in a Nunclon Delta–treated 96-well plate in liquid medium containing 4 % lactose, 2 % spent grain extract, 100 mM PIPPS di-ammonium citrate, 2.4 mM MgSO₄, 4.1 mM CaCl₂ (pH 4.8). The medium was centrifuged and sterile filtered prior to use to reduce the number of particles from the medium during imaging. Spore suspensions were diluted to a final concentration of 1 × 10⁵ spores/mL, resulting in 1 × 10⁴ spores per well in a total well volume of 100 µL. Each strain was grown in 8 replicate wells.

Prior to starting the experiment, inoculated plate was left in the oCelloScope for 2 h to allow spores to settle and focus to be adjusted. Following a 9-hour delay before image acquisition, images were acquired every two hours for a duration of 64 hours using a reduced data acquisition mode. The plate was incubated in the oCelloScope using the integrated BioSense Solutions Mini Incubator during image acquisition. Acquired images were analyzed in UniExplorer software (version 14.3.0.437) using Growth Kinetics (SESAfungi normalized algorithm), Fungal Spore Quantification and Fungi Total Growth analysis modules. Data were extracted from UniExplorer and lag phase duration, exponential growth rate, time to 50 % germination and area under the curve (AUC) were calculated from the data.

Data were analyzed in R v4.5.1 (51). Growth rates of different strains were compared by fitting linear regressions to the exponential phase of SESAfungi normalized growth curves (log_10_ transformed covered surface area) using fixed-size sliding windows (8 time points = 16 hours). The slope of the fitted line was used as a relative measure of growth rate in the exponential growth phase. Lag phase duration was estimated from the x-intercept of the corresponding regression. Germination was expressed as the relative decrease in normalized spore counts over time. Time at 50 % germination was estimated by linear interpolation between consecutive time points. AUC was calculated from baseline-corrected fungi total growth data over the full duration of oCelloScope imaging using the *auc* function from *gcplyr* package (52). Slope, lag phase, time at 50 % germination and AUC estimates were compared between strains using Welch’s one-way ANOVA, followed by Games-Howell post hoc tests, to account for unequal variances. Statistical analyses were performed in R, with Welch’s ANOVA implemented using base R and post hoc tests using the *rstatix* package (53). Significance threshold of *p* < 0.01 was applied.

Noisy segments in the bioreactor time-series were identified and smoothed using generalized additive models (GAMs) with penalized regression splines, fitted using restricted maximum likelihood as implemented in the *mgcv* package in R (54). Gaps arising from removed or unreliable measurements were subsequently filled using robust linear regression MASS package(55), which down-weights the influence of atypical observations during estimation. Savitzky–Golay smoothing was applied using *signal* package function to reduce noise in the carbon evolution rate (CER) time-series data (56).

### SLP2 purification and inhibitor assay

SLP2 production and empty parental strains were grown in 24 deep well plates in 3.5 ml of 4 % glucose, 1 % yeast extract, 15g/l KH_2_PO_4_, 5.4 g/l Na_2_SO_4_, trace elements, 100 mM PIPPS di-ammonium citrate, 2.4 mM MgSO₄, 4.1 mM CaCl₂, pH 4.8. Two plates were collected from each strain and mycelia was removed by filtering through glass fiber filters. Culture pH was adjusted to 7.4 and diluted in half with 20 mM sodium phosphate buffer, pH 7.4 supplemented with 500 mM NaCl and filtered through a 0.45 μM PVDF membrane. SLP2 was purified with HisTrap^TM^ Fast flow immobilized metal ion affinity column and ÄKTA start^TM^ protein purification system (Cytiva). The culture filtrate was fed to the column at a flow rate of 3 ml/min and washed with the same buffer containing 20 mM imidazole. SLP2 was eluted from the column using an imidazole gradient of 0 to 500 mM in 40 min. Fractions containing SLP2 were combined based on SDS-PAGE analysis. Similar procedure was followed with empty parental strain and same fractions were combined from empty parental strain to make the control sample. Both samples were dialyzed overnight at +4 °C against 1xPBS, pH 7.3 using Slide-A-Lyzer™ G3 Dialysis Cassettes, 3.5K MWCO (Thermo Scientific, USA).

Synthetic TrI9 peptide (amino acids 82-151) was ordered from GenScript and reconstituted in MQ water in 1 mg/ml concentration according to provided purity information. *In vitro* inhibition assay was performed by measuring protease activity of the purified SLP2 in the presence of the TrI9 peptide as described above, but with 1x PBS pH 6.65 as the assay buffer. Reactions containing PMSF or chymostatin in 1 mM or 0.01 mM final concentration, respectively, were included as controls.

## Acknowledgements

The authors thank Krista Lehtinen, Satoko Nomura, Niklas Laukkanen and Eero Kiviniemi for excellent technical assistance. This project received funding from the Research Council of Finland under grant agreement no 368374, and VTT’s internal funding. We acknowledge CSC -- IT Center for Science, Finland, for computational resources.

## Data availability

All data are available in the main text and Supplementary Information.

## Notes

### Competing Interest Statement

The authors have declared no competing interest.

## References

1. J. R. Cherry, A. L. Fidantsef, Directed evolution of industrial enzymes: an update. Curr Opin Biotechnol 14, 438–443 (2003).

2. C. Snyman, L. W. Theron, B. Divol, Understanding the regulation of extracellular protease gene expression in fungi: a key step towards their biotechnological applications. Appl Microbiol Biotechnol 103, 5517–5532 (2019).

3. S. Chai et al., Building a Versatile Protein Production Platform Using Engineered *Trichoderma reesei*. ACS Synth Biol 11, 486–496 (2022).

4. C. Hu et al., Efficient Production of the Recombinant Human Lactoferrin in an Engineered *Trichoderma reesei*. J Agric Food Chem 73, 32795–32806 (2025).

5. Y. Sun et al., Extracellular protease production regulated by nitrogen and carbon sources in *Trichoderma reesei*. J Basic Microbiol 61, 122–132 (2021).

6. G. Zhang, Y. Zhu, D. Wei, W. Wang, Enhanced production of heterologous proteins by the filamentous fungus *Trichoderma reesei* via disruption of the alkaline serine protease SPW combined with a pH control strategy. Plasmid 71, 16–22 (2014).

7. C. P. Landowski et al., Enabling low cost biopharmaceuticals: high level interferon alpha-2b production in *Trichoderma reesei*. Microb Cell Fact 15, 104 (2016).

8. C. P. Landowski et al., Enabling Low Cost Biopharmaceuticals: A Systematic Approach to Delete Proteases from a Well-Known Protein Production Host *Trichoderma reesei*. Plos One 10, e0134723 (2015).

9. Y. Qian et al., Enhancement of Cellulase Production in *Trichoderma reesei* via Disruption of Multiple Protease Genes Identified by Comparative Secretomics. Front Microbiol 10, 2784 (2019).

10. C. Yao et al., Overexpression of a Novel Vacuolar Serine Protease-Encoding Gene (*spt1*) to Enhance Cellulase Production in *Trichoderma reesei*. Fermentation-Basel 9, 191 (2023).

11. I. V. Demidyuk, A. V. Shubin, E. V. Gasanov, S. V. Kostrov, Propeptides as modulators of functional activity of proteases. Biomol Concepts 1, 305–322 (2010).

12. H. Ikemura, H. Takagi, M. Inouye, Requirement of pro-sequence for the production of active subtilisin E in Escherichia coli. J Biol Chem 262, 7859–7864 (1987).

13. A. R. Khan, M. N. James, Molecular mechanisms for the conversion of zymogens to active proteolytic enzymes. Protein Sci 7, 815–836 (1998).

14. K. Maier, H. Muller, H. Holzer, Purification and molecular characterization of two inhibitors of yeast proteinase B. J Biol Chem 254, 8491–8497 (1979).

15. P. Schu, P. Suarez Rendueles, D. H. Wolf, The proteinase yscB inhibitor (PB12) gene of yeast and studies on the function of its protein product. Eur J Biochem 197, 1–7 (1991).

16. S. Kojima, Y. Hisano, K. Miura, Alteration of inhibitory properties of *Pleurotus ostreatus* proteinase A inhibitor 1 by mutation of its C-terminal region. Biochem Biophys Res Commun 281, 1271–1276 (2001).

17. M. Hohl, A. Stintzi, A. Schaller, A novel subtilase inhibitor in plants shows structural and functional similarities to protease propeptides. J Biol Chem 292, 6389–6401 (2017).

18. C. Y. Lee et al., Systematic discovery of protein interaction interfaces using AlphaFold and experimental validation. Mol Syst Biol 20, 75–97 (2024).

19. J. K. Varga, S. Ovchinnikov, O. Schueler-Furman, actifpTM: a refined confidence metric of AlphaFold2 predictions involving flexible regions. Bioinformatics 41 (2025).

20. M. van Kempen et al., Fast and accurate protein structure search with Foldseek. Nat Biotechnol 42, 243–246 (2024).

21. J. Abramson et al., Accurate structure prediction of biomolecular interactions with AlphaFold 3. Nature 630, 493–500 (2024).

22. Y. Li, Z. Hu, F. Jordan, M. Inouye, Functional analysis of the propeptide of subtilisin E as an intramolecular chaperone for protein folding. Refolding and inhibitory abilities of propeptide mutants. J Biol Chem 270, 25127–25132 (1995).

23. U. Shinde, Y. Li, S. Chatterjee, M. Inouye, Folding pathway mediated by an intramolecular chaperone. Proc Natl Acad Sci U S A 90, 6924–6928 (1993).

24. R. F. Alford et al., The Rosetta All-Atom Energy Function for Macromolecular Modeling and Design. J Chem Theory Comput 13, 3031–3048 (2017).

25. N. R. Bennett et al., Improving de novo protein binder design with deep learning. Nat Commun 14, 2625 (2023).

26. M. D. Overath et al., Predicting Experimental Success in De Novo Binder Design: A Meta-Analysis of 3,766 Experimentally Characterised Binders. bioRxiv 10.1101/2025.08.14.670059, 2025.2008.2014.670059 (2025).

27. B. Mechler, H. H. Hirsch, H. Muller, D. H. Wolf, Biogenesis of the yeast lysosome (vacuole): biosynthesis and maturation of proteinase yscB. EMBO J 7, 1705–1710 (1988).

28. V. L. Nebes, E. W. Jones, Activation of the proteinase B precursor of the yeast *Saccharomyces cerevisiae* by autocatalysis and by an internal sequence. J Biol Chem 266, 22851–22857 (1991).

29. R. J. Siezen, J. A. Leunissen, Subtilases: the superfamily of subtilisin-like serine proteases. Protein Sci 6, 501–523 (1997).

30. F. Falkenberg, S. Kohn, M. Bott, J. Bongaerts, P. Siegert, Biochemical characterisation of a novel broad pH spectrum subtilisin from Fictibacillus arsenicus DSM 15822(T). FEBS Open Bio 13, 2035–2046 (2023).

31. S. Kojima, M. Deguchi, K. Miura, Involvement of the C-terminal region of yeast proteinase B inhibitor 2 in its inhibitory action. J Mol Biol 286, 775–785 (1999).

32. S. Kojima, A. Iwahara, Y. Hisano, H. Yanai, Effects of hydrophobic amino acid substitution in Pleurotus ostreatus proteinase A inhibitor 1 on its structure and functions as protease inhibitor and intramolecular chaperone. Protein Eng Des Sel 20, 211–217 (2007).

33. P. Slusarewicz, Z. Xu, K. Seefeld, A. Haas, W. T. Wickner, I2B is a small cytosolic protein that participates in vacuole fusion. Proc Natl Acad Sci U S A 94, 5582–5587 (1997).

34. Z. Xu, A. Mayer, E. Muller, W. Wickner, A heterodimer of thioredoxin and I(B)2 cooperates with Sec18p (NSF) to promote yeast vacuole inheritance. J Cell Biol 136, 299–306 (1997).

35. Z. Xu, K. Sato, W. Wickner, LMA1 binds to vacuoles at Sec18p (NSF), transfers upon ATP hydrolysis to a t-SNARE (Vam3p) complex, and is released during fusion. Cell 93, 1125–1134 (1998).

36. J. A. N. Lehmbeck, H. Udagawa (2008) FUNGAL PEPC INHIBITOR. (NOVOZYMES A/S).

37. M. Steinegger, J. Söding, MMseqs2 enables sensitive protein sequence searching for the analysis of massive data sets. Nat Biotechnol 35, 1026–1028 (2017).

38. R. L. Dunbrack, Jr., Res ipSAE loquunt: What’s wrong with AlphaFold’s ipTM score and how to fix it. bioRxiv 10.1101/2025.02.10.637595 (2025).

39. S. Chaudhury, S. Lyskov, J. J. Gray, PyRosetta: a script-based interface for implementing molecular modeling algorithms using Rosetta. Bioinformatics 26, 689–691 (2010).

40. M. Pacesa et al., One-shot design of functional protein binders with BindCraft. Nature 646, 483–492 (2025).

41. M. D. Tyka et al., Alternate states of proteins revealed by detailed energy landscape mapping. J Mol Biol 405, 607–618 (2011).

42. E. C. Meng et al., UCSF ChimeraX: Tools for structure building and analysis. Protein Sci 32, e4792 (2023).

43. N. Aro et al., Production of bovine beta-lactoglobulin and hen egg ovalbumin by *Trichoderma reesei* using precision fermentation technology and testing of their techno-functional properties. Food Res Int 163, 112131 (2023).

44. M. Saric, L. Eisoldt, V. Doring, T. Scheibel, Interplay of Different Major Ampullate Spidroins during Assembly and Implications for Fiber Mechanics. Adv Mater 33, e2006499 (2021).

45. M. Penttilä, H. Nevalainen, M. Rättö, E. Salminen, J. Knowles, A versatile transformation system for the cellulolytic filamentous fungus *Trichoderma reesei*. Gene 61, 155–164 (1987).

46. A. Rantasalo et al., Novel genetic tools that enable highly pure protein production in *Trichoderma reesei*. Sci Rep 9, 5032 (2019).

47. M. Haddad Momeni et al., Discovery of fungal oligosaccharide-oxidising flavo-enzymes with previously unknown substrates, redox-activity profiles and interplay with LPMOs. Nat Commun 12, 2132 (2021).

48. T. D. Schmittgen, K. J. Livak, Analyzing real-time PCR data by the comparative C(T) method. Nat Protoc 3, 1101–1108 (2008).

49. M. G. Steiger, R. L. Mach, A. R. Mach-Aigner, An accurate normalization strategy for RT-qPCR in Hypocrea jecorina (Trichoderma reesei). J Biotechnol 145, 30–37 (2010).

50. T. Jämsä, N. J. Claassens, L. Salusjärvi, A. Nyyssölä, H(2)-driven xylitol production in Cupriavidus necator H16. Microb Cell Fact 23, 345 (2024).

51. R. C. Team, R: A language and environment for statistical computing. MSOR connections 1 (2014).

52. M. Blazanin, gcplyr: an R package for microbial growth curve data analysis. BMC Bioinformatics 25, 232 (2024).

53. A. Kassambara (2020) Pipe-Friendly Framework for Basic Statistical Tests [R package rstatix version 0.6.0].

54. S. Wood, Generalized Additive Models: An Introduction With R*, Second Edition* (Chapman and Hall/CRC, 2017), vol. 66, pp. 391.

55. W. N. Venables, B. D. Ripley, Modern Applied Statistics with S (Springer, 2002), 10.1007/978-0-387-21706-2

56. A. Savitzky, M. J. E. Golay, Smoothing and Differentiation of Data by Simplified Least Squares Procedures. Analytical Chemistry 36, 1627–1639 (1964).

