## Supplemental data for "Protease suppression by native I9 inhibitor improves recombinant protein production in *Trichoderma reesei*"

### Supplementary figures:

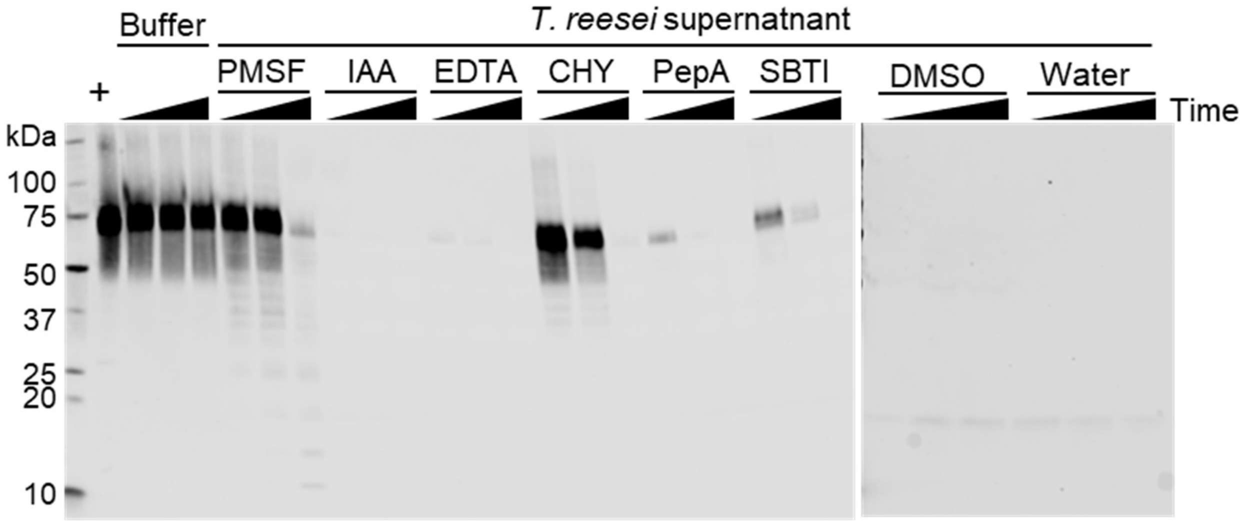

**Fig. S1. CBM-AQ12-CBM is targeted by PMSF- and Chymotrypsin-sensitive proteases in *T. reesei* supernatant.** Purified CBM-AQ12-CBM was incubated without *T. reesei* supernatant (Buffer) or in the presence of *T. reesei* supernatant where different protease inhibitors (PMSF, IAA, EDTA, CHY, PepA, SBTI) or their solvents (DMSO, Water) were added. Protein samples were taken 1.5h, 3.5h and 21h after mixing. The strep-tagged CBM-AQ12-CBM was detected by Western blotting using an Anti-strep-specific antibody. +, 500 ng purified AQ12 loaded.

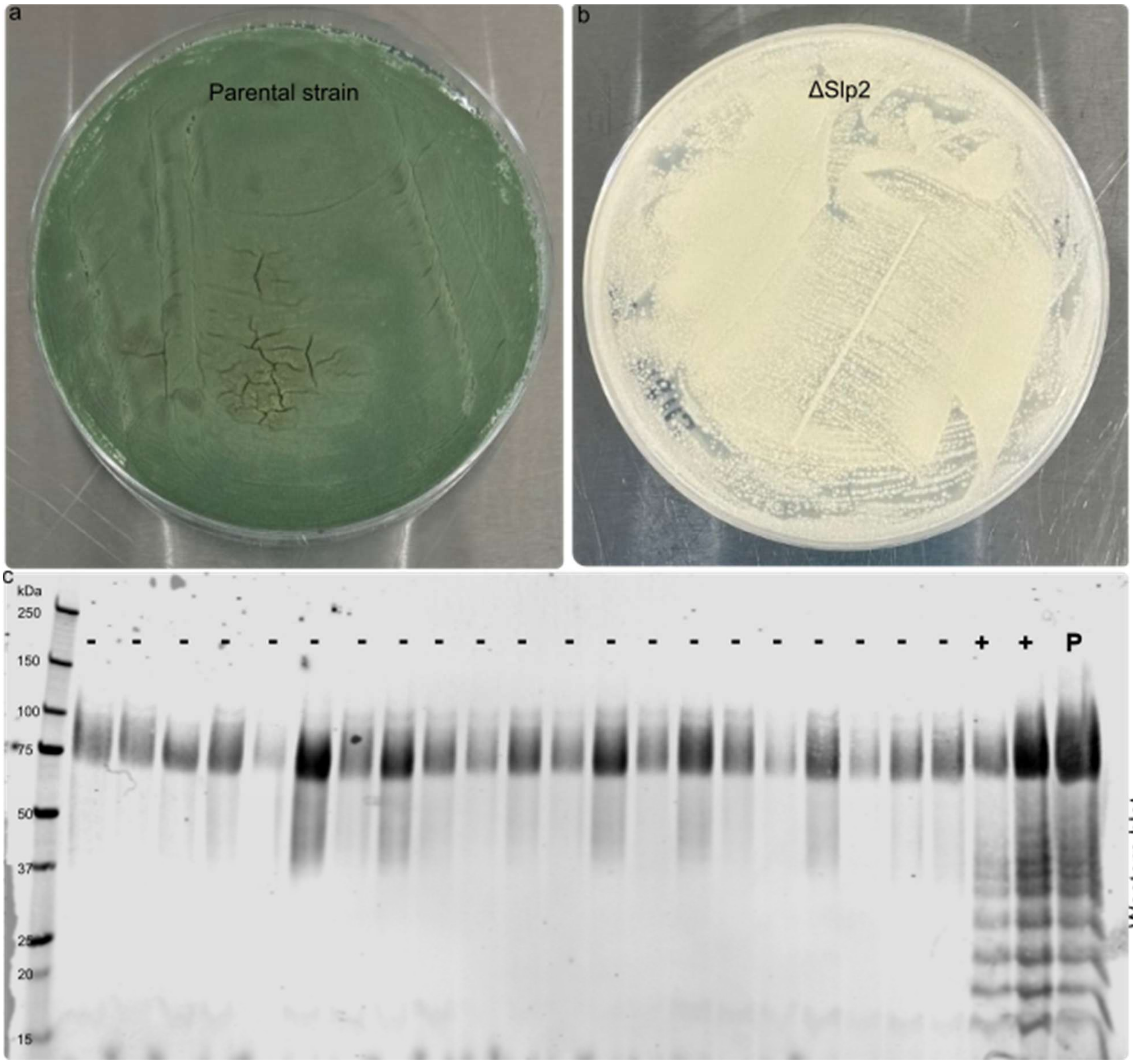

**Fig S2. Deletion of *slp2* prevents normal sporulation of *T. reesei* on PD plates but reduces degradation of CBM-AQ12-CBM.** Parental strain shows normal green sporulation on PD plates (a). Deletion of *slp2* prevents normal spore formation (b). Screening of multiple transformants show that deletion of *slp2* (-) reduces the degradation of CBM-AQ12-CBM compared to parental strain (P) or strain were slp2 deletion was not successful (+). The CBM-AQ12-CBM was detected using antibody against C-terminal Strep II tag.

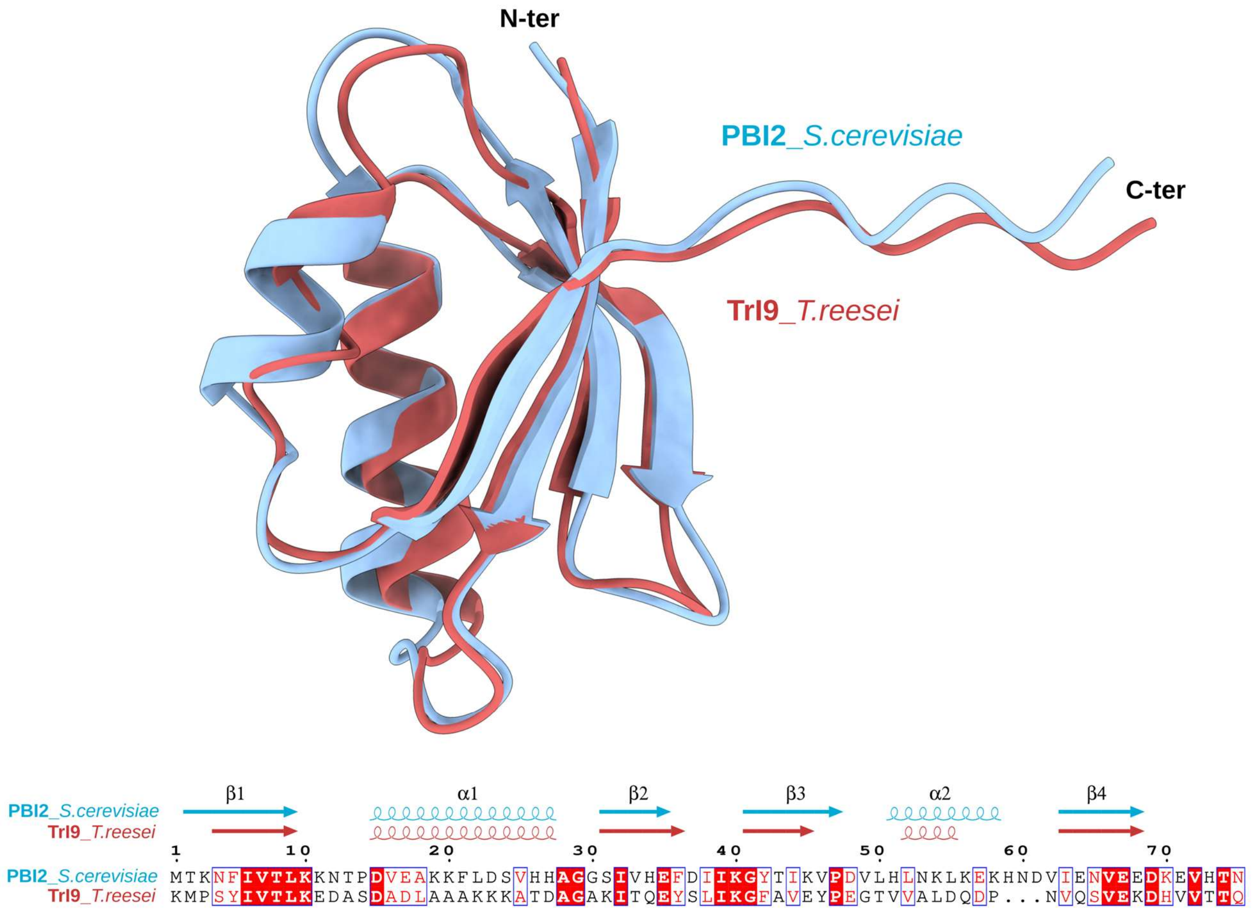

**Fig. S3. Structural conservation between the *S. cerevisiae* Pbi2 inhibitor and the *T. reesei* TrI9 inhibitor.** *(Top)* Superposition of the AlphaFold3-predicted structures of *S. cerevisiae* Pbi2 (blue cartoon, full length) and *T. reesei* TrI9 (red cartoon, residues 81–151; the N-terminal disordered region, residues 1–80, omitted), revealing a conserved I9 fold despite only 28 % sequence identity. *(Bottom)* Structure-based sequence alignment of Pbi2 and TrI9, generated with ESPript 3.2 (Robert & Gouet, 2014). Identical residues are highlighted with red boxes and similar residues with blue boxes. Secondary structure elements derived from the AlphaFold3 models are shown above the alignment as arrows (β-strands) and ribbons (α-helices).

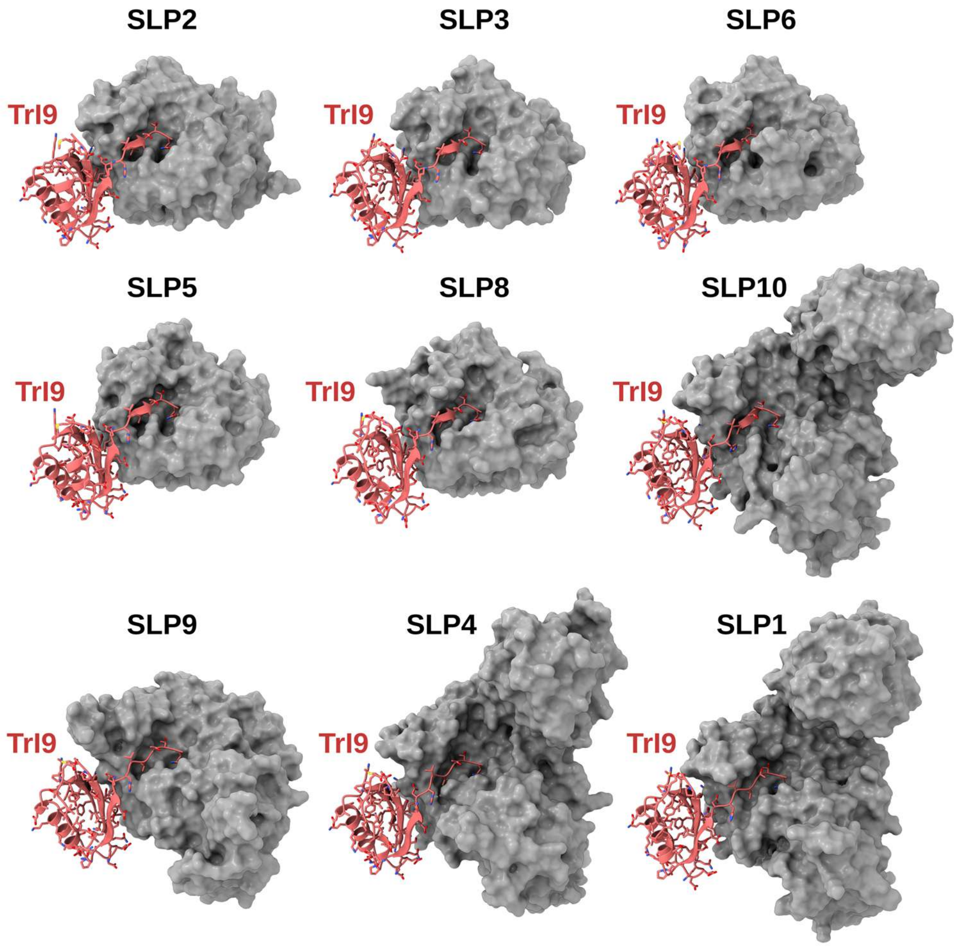

| Binding pair | AF3 ipSAE_min | AF3 interface PAE (Å) | Rosetta interface ΔG (REU) |
|---|---|---|---|
| TrI9 + SLP2 (trire2_123244) | 0.769 | 5.05 | −80.89 |
| TrI9 + SLP3 (trire2_123234) | 0.762 | 4.59 | −85.95 |
| TrI9 + SLP6 (trire2_121495) | 0.739 | 4.96 | −71.38 |
| TrI9 + SLP5 (trire2_64719) | 0.737 | 4.67 | −76.88 |
| TrI9 + SLP8 (trire2_58698) | 0.709 | 5.10 | −81.73 |
| TrI9 + SLP10 (trire2_35726) | 0.198 | 11.92 | −70.31 |
| TrI9 + SLP9 (trire2_60791) | 0.024 | 15.73 | −55.66 |
| TrI9 + SLP4 (trire2_109276) | 0.018 | 16.54 | −67.43 |
| TrI9 + SLP1 (trire2_51365) | 0.016 | 22.19 | −72.22 |

**Fig. S4. Computational prediction of TrI9 binding to *T. reesei* subtilisin-like proteases (SLPs).** *(Top)* AlphaFold3 models of TrI9 (red cartoon with stick side chains) bound to the catalytic domains of nine *T. reesei* SLPs (gray surface), following Rosetta structural relaxation. In all models, the C-terminal loop of TrI9 inserts into the substrate-binding cleft of the protease, with the P1 residue positioned in the catalytic pocket. *(Bottom)* Computational binding metrics for TrI9 complexes with each SLP protease domain, ranked by decreasing AF3 ipSAE_min, an interface-focused AF3 confidence score (higher = more confident) (Overath et al., 2025). Gene identifiers follow the *T. reesei* QM6a genome annotation (trire2). The AF3 interface PAE reports the mean predicted aligned error (in Å), reflecting overall confidence in the predicted binding geometry. Rosetta interface ΔG values (in Rosetta Energy Units, REU) reflect the predicted energetic stability of the complex interface (more negative = more favorable) (Alford et al., 2017).

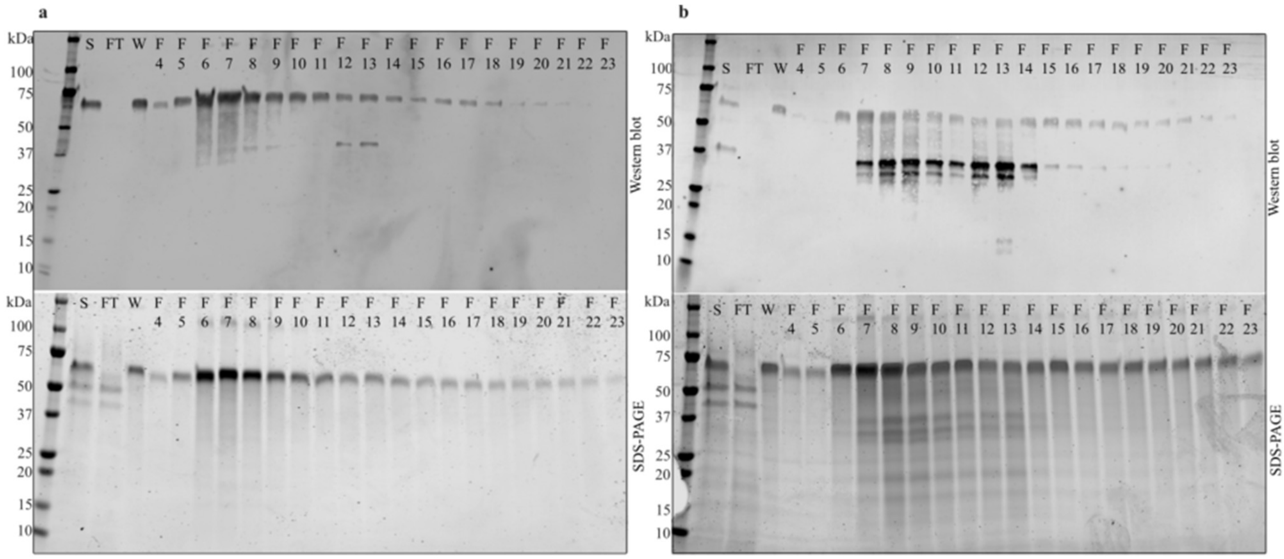

**Fig. S5: Purification of SLP2(12-469-his from *T. reesei* culture supernatants**. Culture supernatants of empty background strain (a) and SLP2 expressing strain (b) was fed to HisTrap$^{TM}$ Fast flow immobilized metal ion affinity column and proteins were eluted with imidazole gradient. Fractions (F) were analyzed with SDS-PAGE and western blot. A double band near the 37 kDa marker was seen in samples from the SLP2(12-469-his) expressing strain (b), but they were absent in similarly treated samples from the background strain (a). Unspecific host cell protein, seen in SDS-PAGE and WB analysis at around the 75 kDa marker was co-purified from both strains. S (input sample), FT (flowthrough), W (wash), F (fractions).

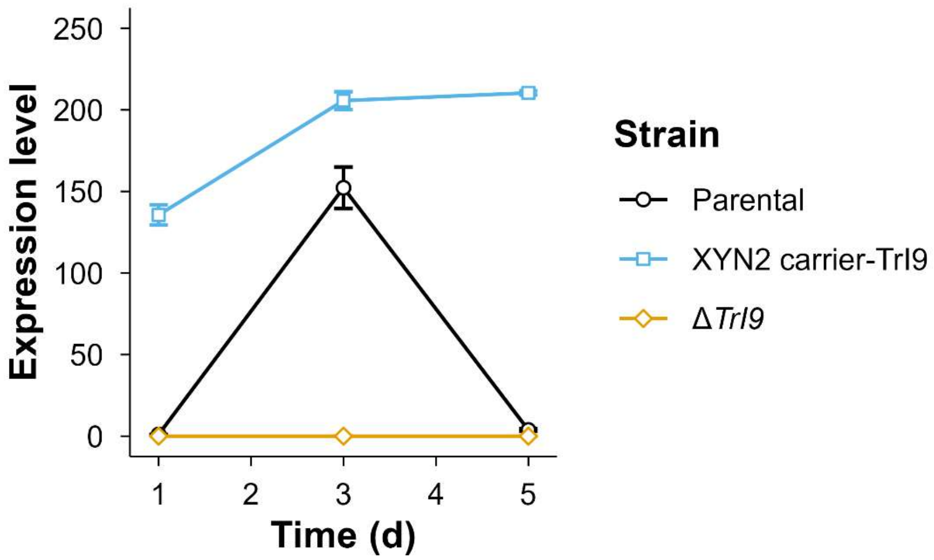

**Fig. S6. Gene expression of TrI9 was analyzed from different strains using RT-qPCR.** The XYN2 carrier-TrI9 showed consistently higher gene expression levels throughout the bioreactor cultivations. *TrI9* deletion strain had essentially no TrI9 expression. Parental strain had upregulated gene expression at day 3 of cultivation after it declined. Expression levels were normalized to *sar1* and *act* expression. Points and error bars represent mean and standard deviation between two qPCR measurements.

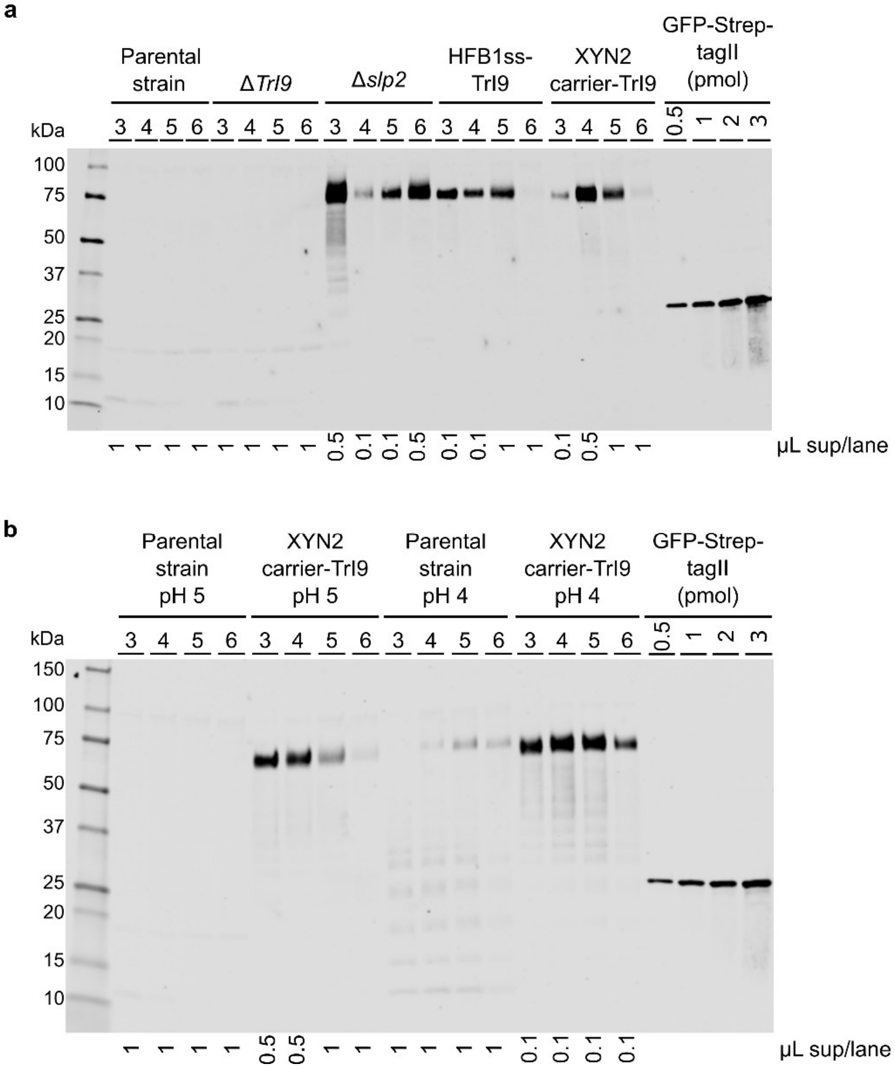

**Fig. S7. Western blots used for the quantification of intact CBM-AQ12-CBM in bioreactor cultivations**. **a** Western blot analysis of culture supernatants from the parental strain, deletion strains (Δ*TrI9* and Δ*slp2*) and *TrI9* overexpression strains (HFB1ss-TrI9 and XYN2 carrier-TrI9) collected on days 3–6. **b** Western blot analysis of culture supernatants from the parental strain and *TrI9* overexpression strain (XYN2 carrier-TrI9) at pH 5 compared to the same strains at pH 4. Supernatants were collected on days 3–6. Detection was performed using a primary antibody against Strep-tag II of CBM-AQ12-CBM. A GFP-Strep-tag II control protein standard curve (0.5–3 pmol, corresponding to 30–180 ng of target protein) was included for quantification. Samples corresponding to 1 µL, 0.5 µL or 0.1 µL of culture supernatant per lane were used for quantification, as indicated below each blot, to ensure signal intensities were in the linear range of the standard curve.

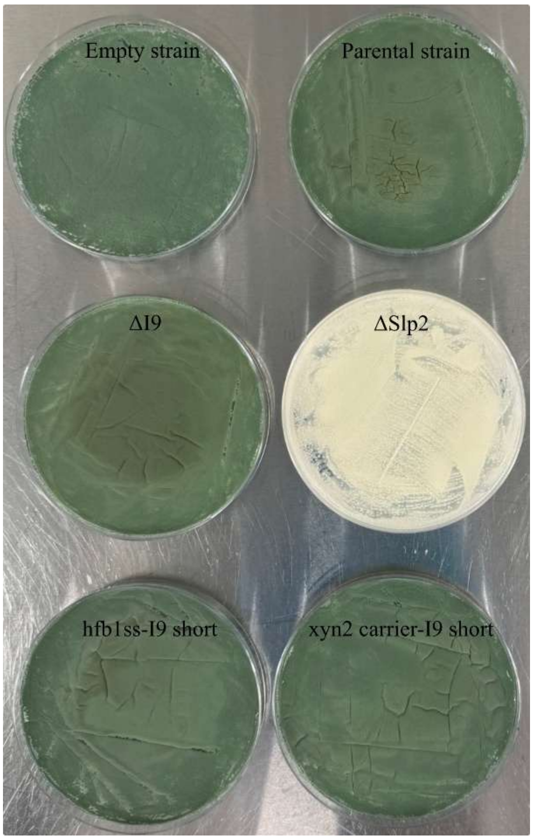

**Fig S8. *Slp2* deletion strain have sporulation defects when grown in PD-agar plates.** The empty background strain, parental stain, TrI9 deletion and TrI9 OE strains sporulated normally.

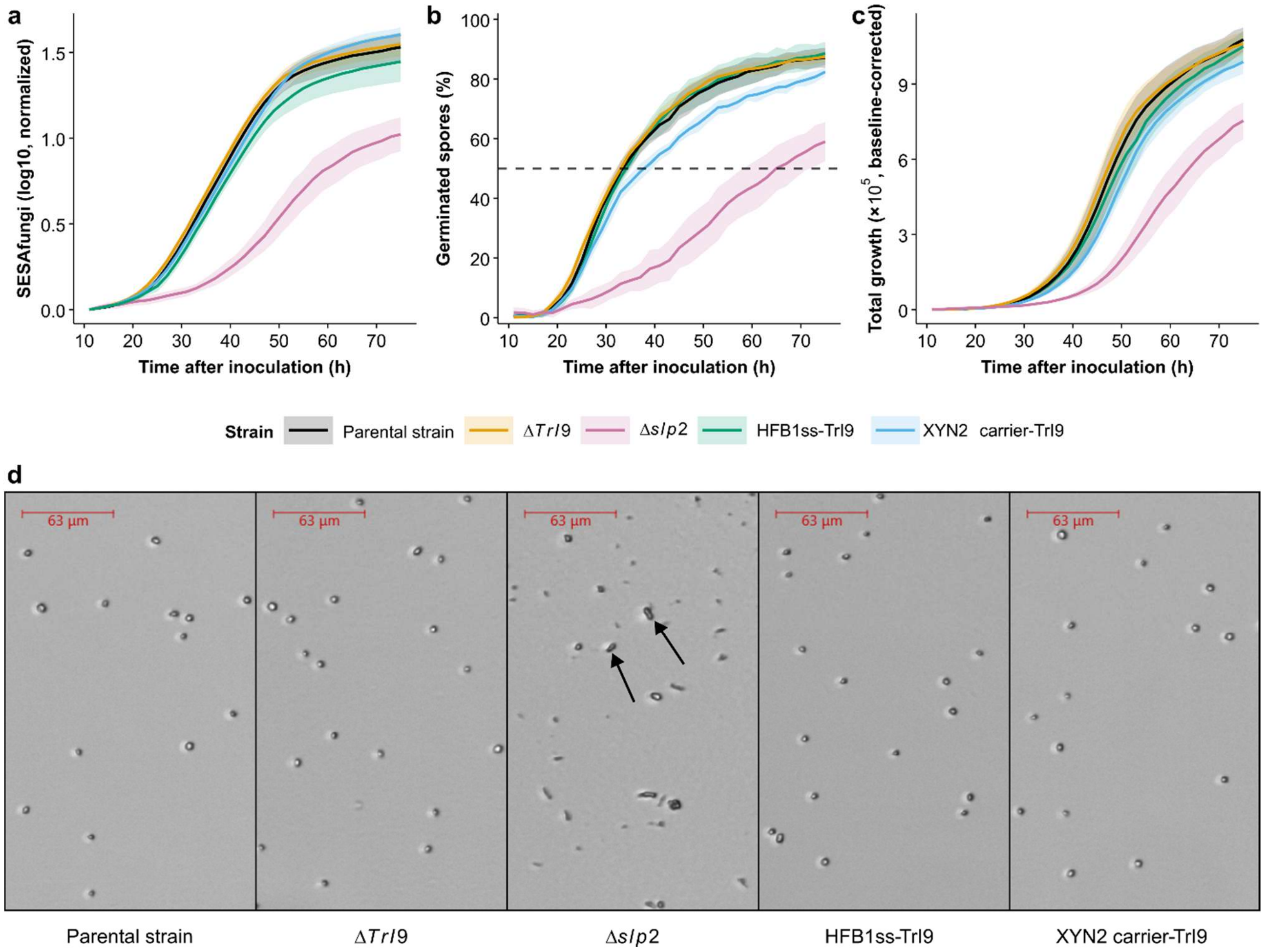

**Fig. S9. Spore germination, growth dynamics and spore morphology of the different *T. reesei* strains. a** Fungal growth kinetics quantified by automated time-lapse microscopy (oCelloScope). Growth curves are based on the SESAfungi normalized signal, reflecting relative changes in fungal surface area over time. **b** Fraction of germinated spores (%) over time measured by automated time-lapse microscopy (oCelloScope). The dashed line indicates the 50 % germination threshold. **c** Total growth of fungal biomass over time measured by automated time-lapse microscopy (oCelloScope). All curves represent the mean of eight replicates per strain and shaded areas around curves represent standard deviation. **d** Microscopy images acquired using oCelloScope showing spores of the different *T. reesei* strains prior to germination (11 h after inoculation). *Slp2* deletion strain has irregularly shaped spores (marked in arrow) compared to round spores in other strains

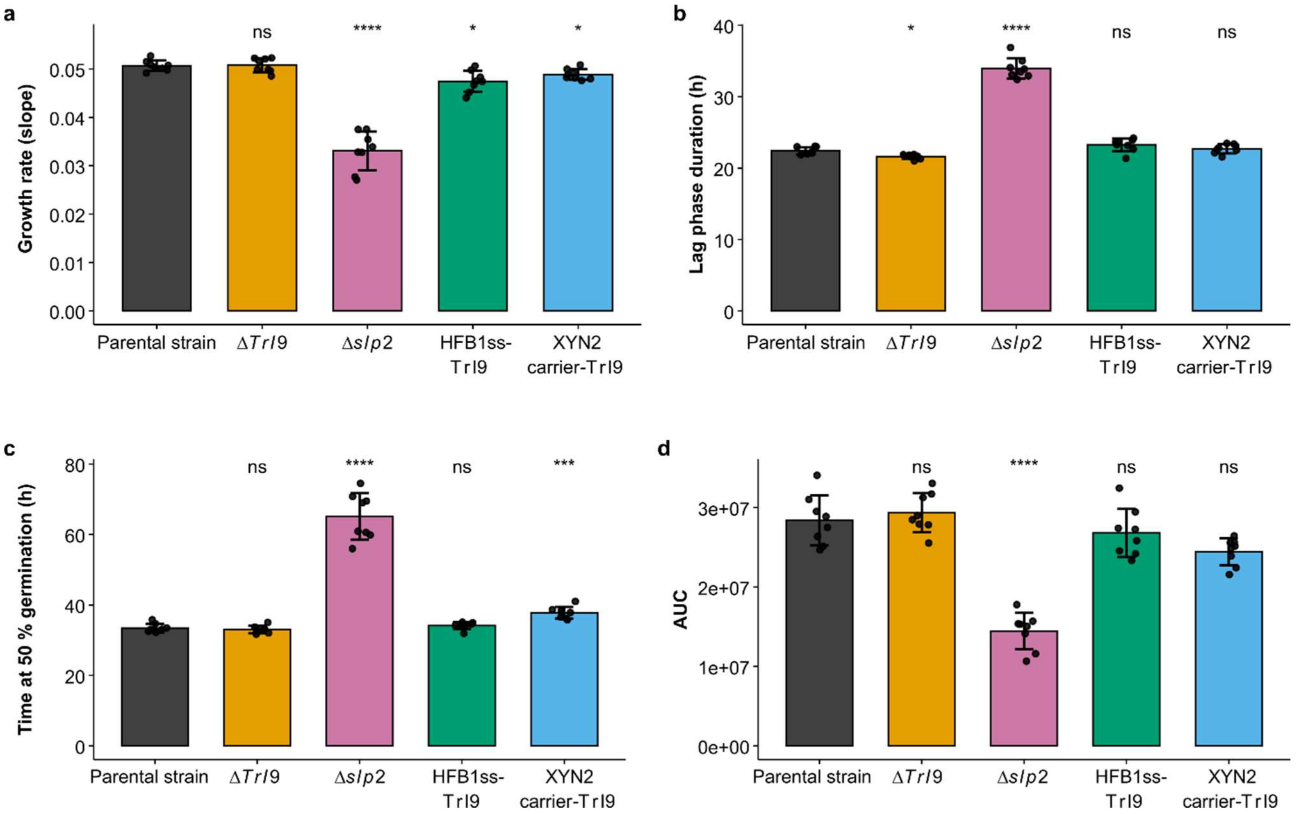

**Fig. S10. Comparison of growth parameters across different *T. reesei* strains.** Growth dynamics of the parental strain and genetically modified strains were quantified using oCelloScope automated time-lapse microscopy. **a** Growth rate was calculated as the slope of a linear fit to SESAfungi normalized data in the exponential growth phase. **b** Lag phase duration, defined as the x-intercept of the fitted slope from SESAfungi normalized data. **c** Time at 50 % germination, calculated from spore quantification data. **d** Area under the curve (AUC), calculated from baseline-corrected fungal total growth data.

All parameters were quantified from microscopy images using oCelloScope image analysis algorithms. The SESAfungi algorithm was used to quantify relative fungal growth based on changes in image-derived fungal surface area over time. The spore quantification algorithm was used to determine germination kinetics based on the decrease in the number of non-germinated spores in microscopy images over time. The fungal total growth algorithm was used to measure the overall fungal biomass accumulation.

Data represent technical replicates (points, n=8) and mean ± SD (bars). Statistical significance was assessed relative to the parental strain using Games-Howell post hoc test, with significance indicated as not significant (ns), $p < 0.05$ (*), $p < 0.01$ (**), $p < 0.001$ (***), $p < 0.0001$ (****).

```
>CBM_AQ12_CMB
HHHHHHHHQACSSVWGQCGGQNWSGPTCCASGSTCVYSNDYYSQCLPGA
STSTGMGPGGGPYGPGASAAAAAAGGYGPGSGQQGPGQQGPGQQGPGQQG
PGQQGPYGPGASAAAAAAGGYGPGSGQQGPGQQGPGQQGPGQQGPGQQGP
YGPGASAAAAAAGGYGPGSGQQGPGQQGPGQQGPGQQGPGQQGPYGPGAS
AAAAAAGGYGPGSGQQGPGQQGPGQQGPGQQGPGQQGPYGPGASAAAAAA
GGYGPGSGQQGPGQQGPGQQGPGQQGPGQQGPYGPGASAAAAAAGGYGPG
SGQQGPGQQGPGQQGPGQQGPGQQGPYGPGASAAAAAAGGYGPGSGQQGP
GQQGPGQQGPGQQGPGQQGPYGPGASAAAAAAGGYGPGSGQQGPGQQGPG
QQGPGQQGPGQQGPYGPGASAAAAAAGGYGPGSGQQGPGQQGPGQQGPGQ
QGPGQQGPYGPGASAAAAAAGGYGPGSGQQGPGQQGPGQQGPGQQGPGQQ
GPYGPGASAAAAAAGGYGPGSGQQGPGQQGPGQQGPGQQGPGQQGPYGPG
ASAAAAAAGGYGPGSGQQGPGQQGPGQQGPGQQGPGQQGGAGGGSGGGQS
HYGQCGGIGYSGPTVCASGTTCQVLNPYYSQCLWSHPQFEK
```

8x-His-tag

Linker sequences

CBHII carbohydrate binding motif (CBM)

12x AQ: PYGPGASAAAAAAGGYGPGSGQQGPGQQGPGQQGPGQQGPGQQG

CBHI CBM

Strep II tag

Theoretical pI/Mw: 6.82 / 57902.51

**Fig. S11 Amino acid sequence and theoretical Mw of CBM-AQ12-CBM.**

### Supplementary tables

**Table S1.** Pre-cultivation cell dry mass

| Strain | cultivation pH | pre-culture biomass g/l |
|---|---|---|
| Parental strain | **pH5** | **2.2** |
| ΔTrI9 | **pH5** | **2.9** |
| Δslp2 | **pH5** | **2.8** |
| hfb1ss-TrI9 short | **pH5** | **3.2** |
| xyn2carrier-TrI9 short | **pH5** | **1.6** |
| Parental strain | **pH4** | **2.2** |
| xyn2carrier-TrI9 short | **pH4** | **2.3** |

**Table S2.** Total protein concentration in bioreactor culture supernatants (g/L)

| Time (d) | Parental | ΔTrI9 | Δslp2 | hfb1ss-TrI9 short | xyn2carrier-I9 short | Parental pH 4 | xyn2carrier-I9 short pl |
|---|---|---|---|---|---|---|---|
| 1 | 2,7 | 2,7 | 2,2 | 2,6 | 2,3 | 0,8 | 1,1 |
| 3 | 5,8 | 6,3 | 3,3 | 5,6 | 4,7 | 4,9 | 5,4 |
| 4 | 9,4 | 9,8 | 4,9 | 7,3 | 6,8 | 8,7 | 7,9 |
| 5 | 13,8 | 11,9 | 7,8 | 11,0 | 9,5 | 12,5 | 15,4 |
| 6 | 14,9 | 16,8 | 11,5 | 15,6 | 15,2 | 19,0 | 21,9 |
| 7 | 20,1 | 19,5 | 14,4 | 17,8 | 16,8 | 23,5 | 25,4 |

**Table S3.** Primers and gRNAs used in the study

| gRNA/oligo | 5'-->3' sequence |
|---|---|
| 5' gRNA TrI9 | CGTATCGATACCTGCCCCGG |
| 3' gRNA TrI9 | TTGTGACGACGCAGTAAACG |
| 5' gRNA slp2 | CGTTGTCGCCCTCTCCATGG |
| 3' gRNA slp2 | GAAGATCCACGATCTCGTCG |
| 5' sar1 oligo | TGGATCGTCAACTGGTTCTACGA |
| 3' sar1 oligo | GCATGTGTAGCAACGTGGTCTTT |
| 5' actin oligo | TGAGAGCGGTGGTATCCACG |
| 3' actin oligo | GGTACCACCAGACATGACAATGTTG |
| 5' TrI9 oligo | CGTCCTACATCGTCACCCTC |
| 3' TrI9 oligo | CACAACGTGGTCCTTCTCGA |

**Table S4.** Growth and germination metrics quantified from the oCelloScope data. Exponential growth rate (slope), lag time duration, time to 50 % germination, and total growth (area under the curve, AUC) are shown as mean ± SD from n = 8 replicates.

| Strain | Slope ($h^{-1}$) | Lag time duration (h) | Time to 50 % germination (h) | AUC (×$10^6$ AU) | Sample size (n) |
|---|---|---|---|---|---|
| Parental strain | 0.051 ± 0.001 | 22.4 ± 0.51 | 33.4 ± 1.24 | 28.4 ± 3.16 | 8 |
| Δ*slp2* | 0.033 ± 0.004 | 34.0 ± 1.42 | 65.1 ± 6.59 | 14.4 ± 2.30 | 8 |
| Δ*TrI9* | 0.051 ± 0.001 | 21.6 ± 0.31 | 33.0 ± 1.05 | 29.3 ± 2.47 | 8 |
| XYN2 carrier-TrI9 | 0.049 ± 0.001 | 22.7 ± 0.63 | 37.8 ± 1.67 | 24.5 ± 1.69 | 8 |
| HFB1ss-TrI9 | 0.047 ± 0.002 | 23.2 ± 0.90 | 34.2 ± 0.99 | 26.8 ± 3.03 | 8 |

**Table S5.** Results of Welch one-way ANOVA for growth and germination parameters derived from the oCelloScope data. The *F* statistics, numerator and denominator degrees of freedom (df) and *p*-values are shown (n = 8).

| Parameter | *F* | df (num, denom) | *p*-value |
|---|---|---|---|
| Slope | 35.89 | 4, 17.04 | $4.3 \times 10^{-8}$ |
| Lag time | 130.98 | 4, 16.55 | $2.7 \times 10^{-12}$ |
| Time to 50 % germination | 50.55 | 4, 17.10 | $3.0 \times 10^{-9}$ |
| AUC | 44.38 | 4, 17.24 | $7.4 \times 10^{-9}$ |

**Table S6.** Adjusted *p*-values for post hoc comparisons to the parental strain using the Games-Howell test for the different growth and germination metrics quantified from the oCellosScope data (n = 8).

| Strain | Slope | Lag time | Time to 50 % germination | AUC |
|---|---|---|---|---|
| Δ*slp2* | $1.5 \times 10^{-5}$ | $3.9 \times 10^{-8}$ | $1.2 \times 10^{-5}$ | $1.6 \times 10^{-6}$ |
| Δ*TrI9* | 1 | 0.020 | 0.97 | 0.96 |
| XYN2 carrier-TrI9 | 0.034 | 0.86 | $4.0 \times 10^{-4}$ | 0.063 |
| HFB1ss-TrI9 | 0.025 | 0.23 | 0.67 | 0.84 |
